# Identification of determinants of differential chromatin accessibility through a massively parallel genome-integrated reporter assay

**DOI:** 10.1101/2020.03.02.973396

**Authors:** Jennifer Hammelman, Konstantin Krismer, Budhaditya Banerjee, David K Gifford, Richard Sherwood

## Abstract

A key mechanism in cellular regulation is the ability of the transcriptional machinery to physically access DNA. Pioneer transcription factors interact with DNA to open chromatin, which subsequently enables changes to gene expression during development, disease, or as a response to environmental stimuli. However, the regulation of DNA accessibility via the recruitment of transcription factors is difficult to understand in the context of the native genome because every genomic site is distinct in multiple ways. Here we introduce the Multiplexed Integrated Accessibility Assay (MIAA), a multiplexed parallel reporter assay which measures changes to genome accessibility as a result of the integration of synthetic oligonucleotide phrase libraries into a controlled, natively inaccessible genomic context. We apply MIAA to measure the effects of sequence motifs on cell type-specific DNA accessibility between mouse embryonic stem cells and embryonic stem cell-derived definitive endoderm cells, screening a total of 7,905 distinct phrases. MIAA is able to recapitulate differential accessibility patterns of 100-nt sequences derived from natively differential genomic regions, identifying the presence of E-box motifs common to epithelial-mesenchymal transition driver transcription factors in stem cell-specific accessible regions that become repressed during differentiation to endoderm. We further present causal evidence that a single binding motif for a key regulatory transcription factor is sufficient to open chromatin, and classify sets of stem cell-specific, endoderm-specific, and shared pioneer factor motifs. We also demonstrate that over-expression of two definitive endoderm transcription factors, Brachyury and FoxA2, results in changes to accessibility in phrases containing their respective DNA-binding motifs. Finally, we use MIAA results to explore the order of motif interactions and identify preferential motif ordering arrangements that appear to have an effect on accessibility.

## Introduction

Genomic DNA acts as an instruction book for the cellular machinery to carry out functional processes such as RNA production^1,2^ and DNA repair^3^. Some regions of the genome are constitutively used across all cell types and states for shared housekeeping processes^4,5^, while other regions are required only in specific cell types as they encode functions that relate to specialized cell properties^6,7^. One key mechanism used by cells to control which regulatory regions can be used in a given cell type is the regulation of chromatin accessibility. Because many transcription factors are incapable of binding in inaccessible or “closed” chromatin, the regulation of chromatin accessibility ensures that such transcription factors do not bind to extraneous or deleterious locations in the genome. In fact, the vast majority of transcription factors bind to <10% of their strong genomic motifs in a given cell type.

“Pioneer” transcription factors are thought to establish the accessibility of cell type-specific regions and initiate cell state change in differentiation^1,8^, cancer^9,10^, and environmental responses^11,12^ and allow “settler” transcription factors to bind and activate previously inactive genes. Massively Parallel Reporter Assays^13,14^ have been developed to measure the change to gene expression from the action of promoters^15,16^ or enhancers^17–22^, and thus can be used to probe the regulatory code either exogenous to the genome via transiently transfected plasmids^17,19^, or integrated into the genome via lentiviral infection^20,21,23^. One area of study made easier by the advent of MPRAs is understanding the combinatorial logic of transcription factor action, such as whether specific combinations of transcription factor binding sites must be co-localized to cause specific genes to be expressed as a hypothesized “enhanceosome” model of genomic regulation^18,24,25^, or whether non-specific additive activities of transcription factors is sufficient^26,27^. However, MPRAs do not measure changes to chromatin accessibility and thus cannot disentangle gene regulation by transcription factors that depend upon changes in local accessibility.

Previous work has indicated specific transcription factor motifs and logic governing chromatin accessibility^28–30^, but such effects are difficult to disentangle in a native genomic context, where motifs are not independent of non-local sequence effects. Recent approaches have extended MPRAs to measure nucleosome occupancy via bisulfite treatment^31^ or MNase-seq^32^ in yeast. However, bisulfite sequencing requires constrained library design to ensure sufficient CpG sites which act as a substrate for bisulfite conversion, and MNase-seq requires measurement over multiple MNase concentrations to fully measure accessibility^33^. Here, we introduce an assay for measuring the local DNA accessibility of genomically integrated large-scale reporter libraries. We demonstrate with this Multiplexed Integrated Accessibility Assay (MIAA) that we can identify DNA sequence motifs which drive cell type-specific changes to accessibility, and that some of these changes can be explained by the expression of cell type-specific transcription factors. Finally, we observe interactions between transcription factors that point to an effect of preferential motif arrangement on accessibility.

## Results

### I Multiplexed Integrated Accessibility Assay (MIAA) measures local accessibility of integrated phrases

In previous work, we used a DNase I cleavage assay to measure chromatin accessibility of a set of phrases integrated into a defined genomic locus^27^. While effective at determining the relative accessibility of classes of DNA sequences, this assay did not allow us to obtain reproducible estimates of chromatin accessibility for individual DNA sequences in the context of high-throughput libraries. We hypothesized that we could measure changes in DNA accessibility at a closed locus with higher sensitivity by observing the conditional binding of an inert adenine DNA methyltransferase to the locus, given the substantially higher efficiency of Dam methylation as compared to DNase digestion of native chromatin (unpublished observations).

We first integrated a set of 100-nt synthesized oligonucleotide phrases that contain a GATC restriction site at a natively closed genomic locus proximal to a retinoic acid receptor (RAR) binding site (Figure 1a). After phrase integration, we expressed a novel protein construct comprising a settler transcription factor, RAR, fused to a mutant version of Dam methyltransferase shown to display increased signal-to-noise over wild-type Dam^34,35^. We expected that phrases that open chromatin should result in increased adenine methylation of the phrase’s GATC site due to the combined effect of increased local RAR binding and the known preference of Dam methylase to methylate in accessible chromatin^35^. We then extracted genomic DNA from the cells, split it into two pools, and exposed one pool to the restriction enzyme DpnI and the other pool to DpnII which preferentially cleave methylated and unmethylated GATC sites, respectively. In each pool, we produced PCR amplicons that crossed the GATC site and included the adjacent phrase, using nextgen sequencing (NGS) of the two pools to produce two count measurements for each phrase (Figure 1b). The PCR primers were designed to exclusively amplify phrases that were correctly integrated into the expected genomic site to avoid amplification of unintegrated or erroneously integrated phrases. The proportion of DpnI to DpnII sequencing counts represents the impact of that phrase on local DNA accessibility (Figure 1c). We designate this high-throughput genomically integrated assay of chromatin accessibility the Multiplexed Integrated Accessibility Assay (MIAA). Since our particular interest was in the changes to accessibility during differentiation, we measured our phrase library in mouse embryonic stem cells (stem cells) as well as mouse embryonic stem cell-derived definitive endoderm cells (definitive endoderm) produced using a well-established differentiation protocol shown to yield >90% definitive endoderm^36^.

**Figure 1.**
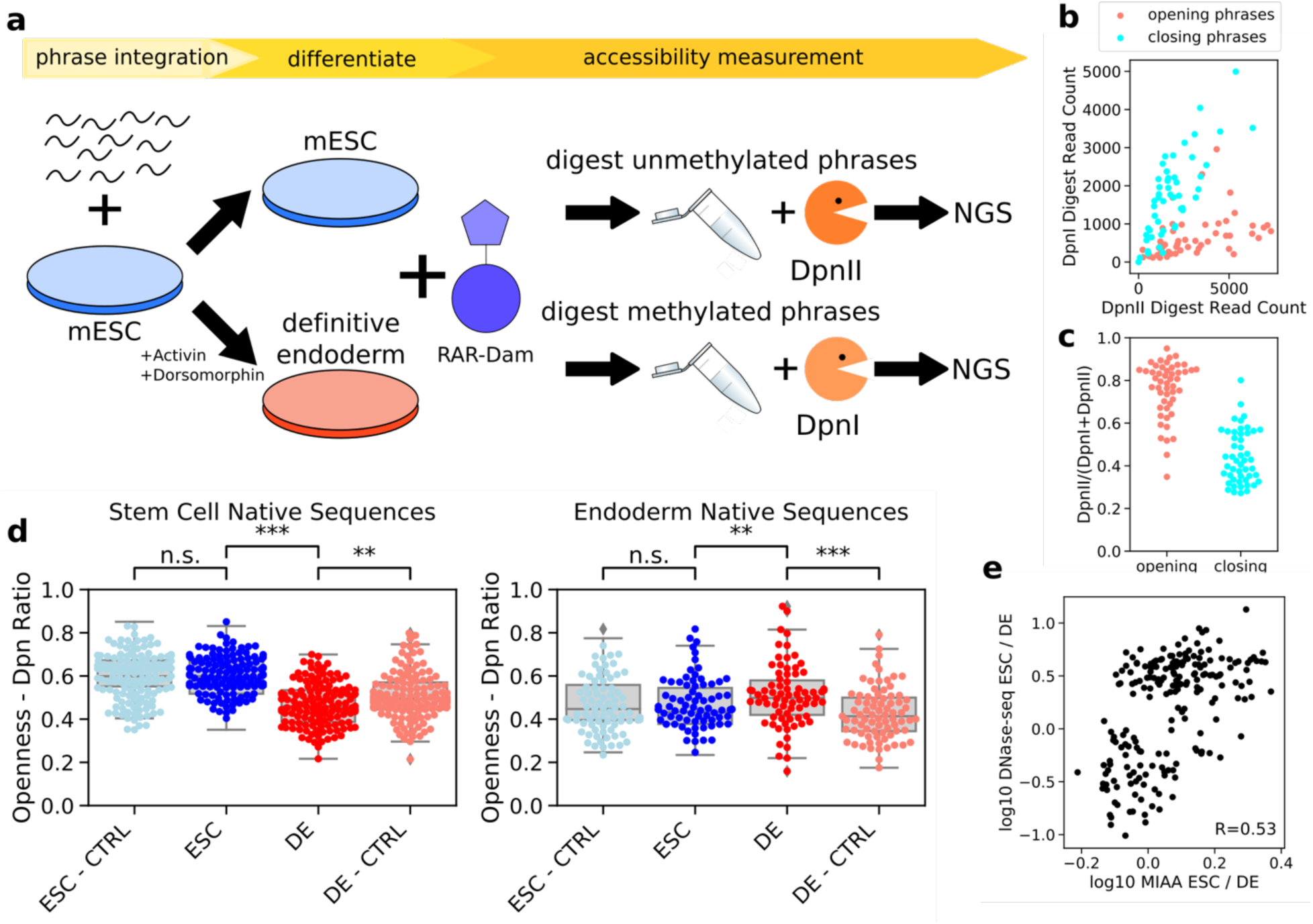
Multiplexed Integrated Accessibility Assay (MIAA) measures local DNA accessibility of synthesized oligonucleotide phrase libraries. A) 100nt phrases are integrated into stem cells at a designated genomic locus. Stem cells are split and half are differentiated into definitive endoderm cells. Retinoic acid receptor fused to hyper-activated deoxyadenosine methylase (DAM) enzyme results in methylation of phrases that open DNA. DNA is extracted and half is exposed to DpnII, which cleaves unmethylated sequences, while half is exposed to DpnI, which cleaves methylated sequences. Sequences are PCR amplified and sequenced. B) DpnI and DpnII read counts measured from a single definitive endoderm replicate show difference between designed opening and closing phrases. C) Proportion of DpnII read counts measured from a single definitive endoderm replicate gives estimate of MIAA openness. D) 100nt native sequences that were differentially opening as measured by DNase-seq are differentially opening as measured by MIAA and different from randomly shuffled control sequences (significance measured by paired t-test). E) Differential accessibility as measured by log change in normalized DNase-seq reads and MIAA methylation proportion shows correlation between native differential accessibility and MIAA accessibility. The correlation reported is the Pearson correlation coefficient (r).

We first tested a library of 5,978 phrases in 4 biological replicate phrase integrations for stem cell and 2 biological replicate phrase integrations for endoderm. To gauge the reliability of MIAA, we included sets of positive and negative control phrases, sampled from the universally opening and universally closing phrases generated and tested in Hashimoto et al^27^. Results from cell type-specific replicates were consistent (Pearson’s r = 0.509-0.848; Supplementary Figure 1). We found that the positive control phrases yielded significantly higher Dam methylation than the negative control phrases (Figure 1b-c), with an average of 87/100 positive control phrases yielding higher methylation than the average negative control phrase in each replicate (p < 0.001 by Wilcoxon rank-sum test for all replicates), suggesting that MIAA can resolve accessibility induction of single phrases in the context of large libraries.

We then examined 213 native genomic sequences that were included in this library. These native sequences were predicted to be differentially accessible between stem cells and definitive endoderm cells by a deep learning model trained to predict DNase-accessible regions from underlying DNA sequence. This method, which we call DeepAccess, trains an ensemble of 10 convolutional neural networks on DNase-seq data from stem cells and definitive endoderm to predict whether a 100nt genomic region is accessible or inaccessible in both cell types (see Methods for details). We also included in our library a randomly shuffled control counterpart for each phrase in order to account for any potential effects of nucleotide composition. Indeed, these native DNA phrases were found to be differentially accessible in stem cells and endoderm (Figure 1d) with an effect size that correlates with differential accessibility measured by DNase-seq (Pearson’s r=0.53; Figure 1e). While statistically significant as a group, only a subset of the native genomic phrases recapitulated the differential accessibility of the native loci from which they were derived, which may result from the 100-nt phrase not containing all of the binding elements controlling accessibility of the native locus.

Against our expectations, we found that rather than being opening in stem cell compared to shuffled controls, a majority of stem cell native phrases were differentially closing in endoderm compared to shuffled control phrases (Figure 1d). While observing repressed accessibility of native phrases compared to the accessibility of random DNA integrated into a genomic locus that is expected to be endogenously closed is surprising, it is possible that the locus becomes more open during phrase integration as has been observed in other Cas9 experiments^37^ or the locus becomes packed into heterochromatin due to active repression^38^. We performed motif enrichment (see Supplementary Methods for details) on these phrases and found that 98% contained a match to the Zeb2 motif, a known transcriptional repressor that has been implicated in early gastrulation by repression of E-cadherins^39^, as compared to 0% of endoderm native sequences. In contrast, none of our DeepAccess-selected native genomic phrases contained motifs for the known stem cell pioneer factors Oct4, Sox2, or Klf4^8^, which would be predicted to open chromatin in stem cells.

To investigate why DeepAccess chose stem cell native genomic sequences that contain Zeb2 motifs over known pioneer factors, we compared DeepAccess-predicted differential accessibility effect size for consensus motifs for the known pluripotency factors Oct4, Sox2, and Klf4 along with Zeb2 and found that Zeb2 motifs had the strongest predicted effect on differential accessibility (Supplementary Figure 2). Zeb2 binding sites were also enriched in stem cell-specific genomic accessible regions with 14% containing a Zeb2 motif relative to 9% in endoderm-specific accessible regions (p < 0.001 by hypergeometric test). In comparison, 12% of genomic stem cell-specific accessible regions contained a Sox2 site, 6% contained Oct4, and 6% contained Klf4. KEGG biological pathway analysis of Zeb2 motif sites in stem cell accessible regions showed an enrichment of motif sites proximal to genes regulating pluripotency of stem cells (p < 0.001), including the key pluripotency regulators Klf4, Sox2, and Nanog, a finding which is consistent with a model of Zeb2 repression of pluripotency during definitive endoderm differentiation^40^. The Zeb2 motif is highly similar to motifs of other E-box epithelial-mesenchymal transition driver TFs such as Zeb1, Snail, Slug, and Twist^41^, all of which are expressed during stem cell differentiation to endoderm. Overall, we find that 100-nt phrases extracted from genomic regions with differential chromatin accessibility recapitulate this differential accessibility when transplanted to a fixed, natively closed chromatin locus.

### II DNase-seq analysis identifies motifs driving cell type-specific accessibility

We then hypothesized that we could identify and confirm with MIAA sequence features that control chromatin accessibly in a cell type-specific manner through a set of synthetic, designed phrases included in our MIAA tested library. Using cell type-specific DNase-seq data, we extracted short (8-12nt) DNA sequence motifs which we hypothesized would cause differential accessibility using two methods (Figure 2a). First, we applied KMAC^42^, a method for detecting motif enrichment, to two sets of cell type-specific accessible sequences. Second, we used DeepAccess to obtain hypotheses about which motifs were likely to be most responsible for differential accessibility between definitive endoderm and stem cells (see Supplementary Methods for details). Unlike KMAC’s pure enrichment approach, DeepAccess is able to learn nonlinear relationships between sequence motifs for predicting accessibility. From our set of motif hypotheses from both methods, we designed synthetic phrases with either 7 instances of 1 motif (Figure 2b) we call *motif phrases* or 2 different motifs (Figure 2c) we call *motif pair phrases* inserted into 24 fixed sequence backgrounds of varied GC-content, that were previously measured to have a neutral impact on cell type-specific accessibility with MIAA (see Methods for details). For each phrase, we also included a control where the nucleotides are randomly shuffled to observe the influence of nucleotide content alone. We collected MIAA data for this library in 4 replicates in stem cells and 2 replicates in endoderm.

**Figure 2.**
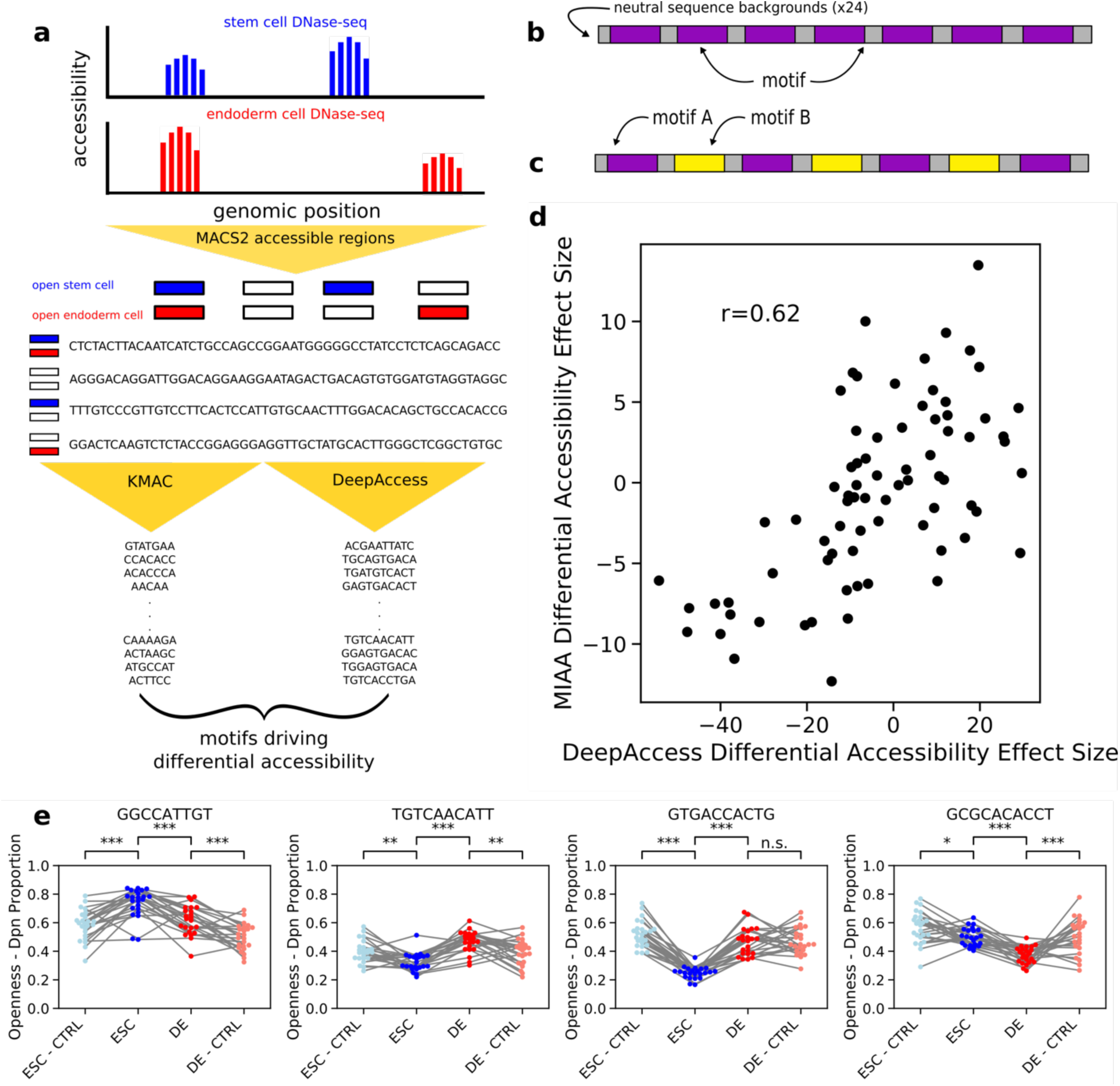
Differentially accessible motif generation from DNase-seq data validated by MIAA. A) DNase-seq accessible regions called with MACS2 and 100nt sequences extracted centered at narrow peak. KMAC and DeepAccess were applied to extract significant motifs potentially driving differential accessibility between stem cells and endoderm. B) Phrases were designed using 7 instances of each motif at the same locations in each phrase inserted into 24 100nt neutral sequence background, as well as C) pairs of motifs. D) Predictions from DeepAccess for differential accessibility replicate experimental results (effect size by paired t-test between stem cell and definitive endoderm measurements). The correlation reported is the Pearson correlation coefficient (r). E) Phrases designed with motifs show differential accessibility via opening stem cell, opening endoderm, closing stem cell, and closing endoderm (left to right). Each dot is an individual phrase and lines are drawn between phrases in each cell context as well as between a phrase and its paired shuffled control (CTRL). Significance computed by paired t-test.

To determine whether DeepAccess was able to predict the effects of computationally selected motif phrases or motif pair phases, we compared the DeepAccess-predicted effect size of each motif or motif pair on differential accessibility to the differential measured effect size with MIAA. We found that DeepAccess results are correlated (Pearson’s r=0.62) with MIAA-measured differential accessibility (Figure 2d). However, we found that DeepAccess failed to perform well in predicting paired effects between phrases and shuffled controls (ESC Pearson’s r=0.24; DE Pearson’s r=0.42; Supplementary Figure 3), which we hypothesize is the result of overconfidence of neural networks on out-of-distribution inputs^43,44^, since the network had not seen entirely random DNA like the shuffled control phrases during training. We tested for statistically significant differential accessibility of our motif hypotheses by first performing paired tests between MIAA openness in stem cell and definitive endoderm and then performing paired tests between phrases and shuffled controls under Benjamini-Hochberg multiple hypothesis correction at a false discovery rate of 0.05 (see Supplementary Methods for details). Out of 38 tested motif phrases, 20 induced differential accessibility, and out of 38 motif pair phrases, 26 induced differential accessibility. We also found these results to be largely consistent across a secondary closed integration locus (Supplementary Figure 4). Thus, MIAA analysis was able to confirm that motifs identified using DeepAccess are able to drive changes to accessibility both between cell types and compared to shuffled controls via local DNA opening or closing (Figure 2e).

Out of the 46 motif or motif pair phrases that induced differential accessibility across cell types and compared to shuffled control phrases as measured by MIAA, DeepAccess predicted the correct direction of differential accessibility between the two cell types in 76% (35/46) of cases (Supplementary Table 1). In comparing results from DeepAccess to KMAC, we found only 32% (8/25) of our KMAC motifs or motif pairs were differentially accessible as compared to 74.5% (38/51) of DeepAccess (Supplementary Table 1), indicating our DeepAccess approach was very successful in identifying accessibility-driving motifs.

### III GC-content and transcription factor binding motifs control accessibility

We noticed previously that the positive control phrases from Hashimoto et al. 2016^27^ had higher GC-content than the negative control phrases. To clarify the role of GC-content in driving accessibility, we included in our library a total of 200 positive and negative control phrases from the Hashimoto et al. 2016 library, which were designed to include a string of motifs correlated with high or low accessibility across cell types^27^. We selected positive and negative controls with either high GC-content (60-70%) or low GC-content (30-50%). We found that in both cell types positive control phrases drove uniformly and equivalently high accessibility regardless of GC-content (Figure 3a), suggesting that motifs associated with accessible regions can increase accessibility independently of GC-content. However, in endoderm, positive control phrases for both GC-content bins had increased accessibility as compared to negative control phrases with matched GC-content (p < 0.001 by Wilcoxon rank-sum test) whereas in stem cells, only the low GC-content bin had differential accessibility between negative and positive controls (p < 0.001 by Wilcoxon rank-sum test) (Figure 3a) because of high accessibility among high-GC negative control phrases. GC-content was positively correlated with openness in both stem cells and definitive endoderm cells among both sets of control phrases (stem cell Pearson’s r=0.476; Pearson’s r=0.357), suggesting that GC-content is a contributor to MIAA-measured accessibility alongside motif composition.

**Figure 3.**
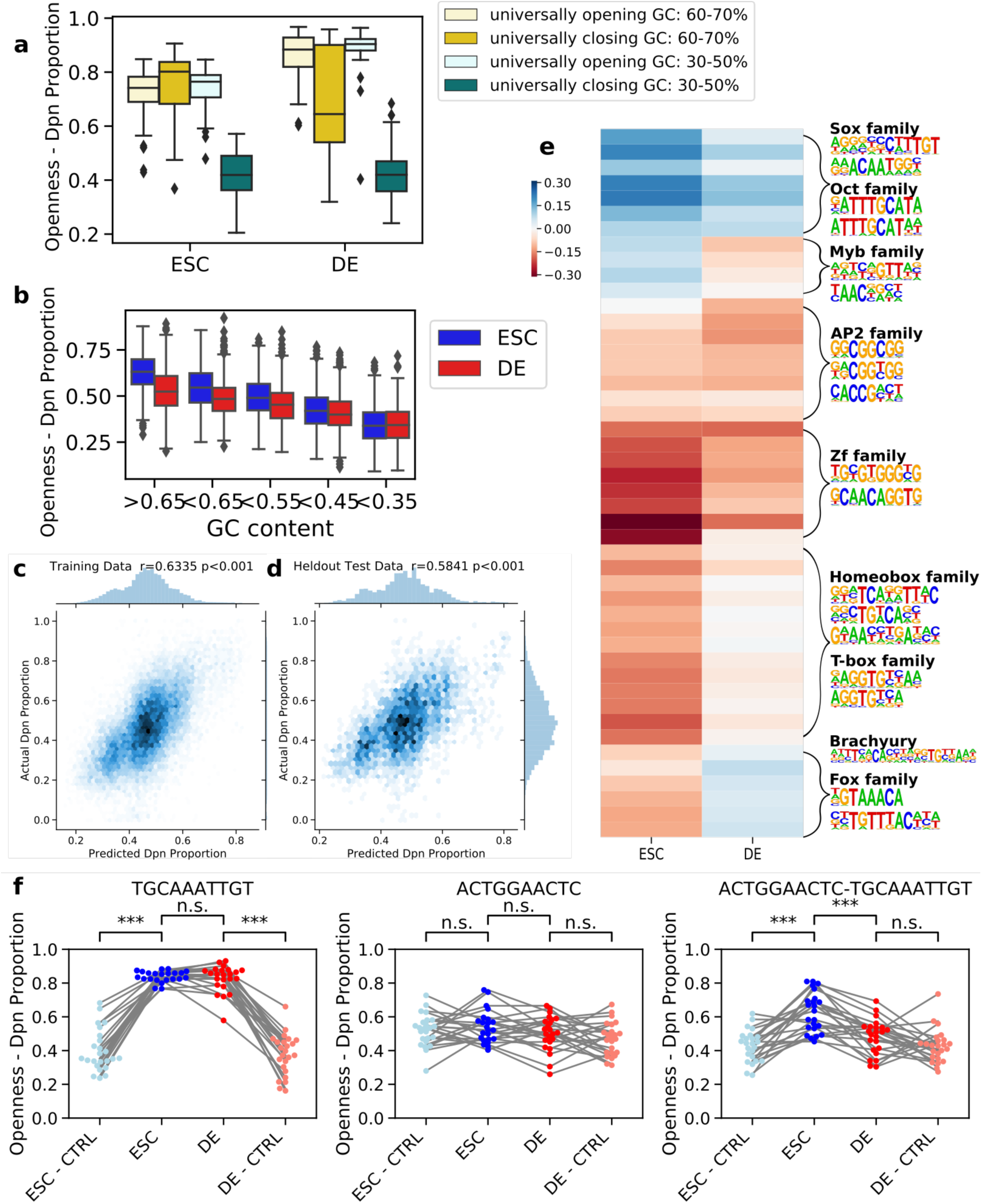
MIAA identifies global influence of GC-content and differentially accessible motifs. A) GC-content observed to be correlated with accessibility in both stem and endoderm cells from positive (universally opening) and negative (universally closing) control sequences. B) GC-content correlated with accessibility in random DNA phrases. The regression model was trained on MIAA Dpn proportions with GC-content, replicate, and cell type-specific effects of 20 motifs and 26 motif pairs as features, and predicts well on (C) training data (n = 21,420) and (D) held-out test data (n = 4,404). The correlation reported is the Pearson correlation coefficient (r). E) Regression weights of individual motifs and motif pairs in stem and definitive endoderm cells. Hierarchical clustering of regression weights followed by motif enrichment recovers clusters representing cell type-specific transcription factor DNA binding motifs. F) Example of individual motifs (left, middle) which alone do not differentially open chromatin, but differentially open chromatin stem cells in combination (right). Each dot represents the average DpnII read proportion of an individual phrase, compared to shuffled controls (CTRL). Significance computed by paired t-test.

Since this result could be an effect of sequence motifs included in the high-GC-content negative control phrases, we then examined the shuffled phrases that we designed to act as controls for motif activity (see previous section) to see if the effect of GC-content on MIAA accessibility held in entirely random DNA. We found that GC-content of randomly shuffled sequences correlated with MIAA accessibility in both cell types (Figure 3b). We also found that accessibility was significantly higher (p < 0.001 by one-tailed Wilcoxon Signed-Rank Test) in stem cells compared to endoderm cells across all GC-content bins (GC-content greater than 0.65, GC-content between 0.55-0.65, GC-content between 0.45-0.55), except in phrases with less than 35% GC-content (N=372). Altogether, these results indicate that GC-content alone is a sufficient DNA signal to drive accessibility in both stem cells and endoderm and also to drive accessibility differences between these two cell contexts through its heightened impact in stem cells.

This finding is consistent with other research that suggests a relationship between GC-rich regions and accessibility^7,33,45^. Consistent with this research, we found that the top 5,000 definitive endoderm-specific regions and the top 5,000 stem cell-specific regions from DNase-seq have higher GC-content than randomly sampled closed regions (Supplementary Figure 5). To estimate the impact each motif or motif pair phrase has on accessibility beyond the effect of GC-content, we then trained a linear regression model to predict MIAA Dpn ratios from GC-content, experimental replicate, and cell type-specific effects for all phrases containing differential motifs or motif pairs. This linear model had good performance on training (Pearson’s r=0.6335; Figure 3c) and held-out test data (Pearson’s r=0.5841; Figure 3d) (see Supplementary Methods for details), and significantly improved from regression models that did not include motif effects, including GC-content and replicate effects alone (adjusted R-squared motif model = 0.398; adjusted R-squared no motif model = 0.095; Supplementary Figure 6), reinforcing the salient effects of transcription factor binding motifs in controlling accessibility. We clustered the regression weights of the motifs and found clusters of motifs representing similar influences on MIAA-measured accessibility. We then ran motif discovery on the designed phrases in each cluster to obtain representative motifs (See Supplementary Methods for details). The identified motifs included known transcription factors such as Oct and Sox motifs in stem cells and T-box and Fox motifs in definitive endoderm (Figure 3e). These regression weights were robust, showing high consistency between models trained on individual biological replicates (Supplementary Figure 7). We also identified motif pair phrases which show interesting non-linear activity with respect to differential accessibility compared to their motif phrase effects alone (Figure 3f). In sum, MIAA data enables *de novo* discovery of features such as GC-content and transcription factor motifs that govern differential chromatin accessibility.

### III Overexpression of definitive endoderm transcription factors Brachyury and FoxA2 increase accessibility of phrases with their DNA-binding motifs

We then hypothesized we could connect our discovered motifs to the causal opening of chromatin by ectopically expressing the transcription factors corresponding to the motifs. We over-expressed the transcription factors Brachyury or FoxA2 in the stem cells and measured the accessibility of our phrase library with MIAA (Figure 4a). We trained a joint regression model to predict condition-specific accessibility with data from four conditions: stem cells, definitive endoderm cells, stem cells with FoxA2 over-expression and stem cells with Brachyury over-expression (see Supplementary Methods for details). We then selected the motifs which had the greatest positive difference in regression weights which indicate an increase in accessibility between the over-expressed Brachyury (ESC Brachyury+) and the stem cell (ESC) conditions, and found that the top effect was a motif pair which partially matches the motif of a Brachyury homodimer with two motifs in a minus/plus orientation and is significantly enriched over other dimer orientations in Brachyury ChIP-seq peaks (p < 0.001 by Chi-squared test; Supplementary Figure 9). The second strongest motif was also significantly enriched in the DNA sequence of ChIP-seq of Brachyury binding in mouse definitive endoderm (p < 0.05 under Benjamini-Hochberg multiple hypothesis correction; Figure 4b). Overall, only 6/76 motifs or motif pairs showed a significant increase in accessibility upon Brachyury over-expression compared to stem cells using our method for measuring statistically significant differential accessibility (see Supplementary Methods for details), showing the specific effect of such over-expression on Brachyury motifs.

**Figure 4.**
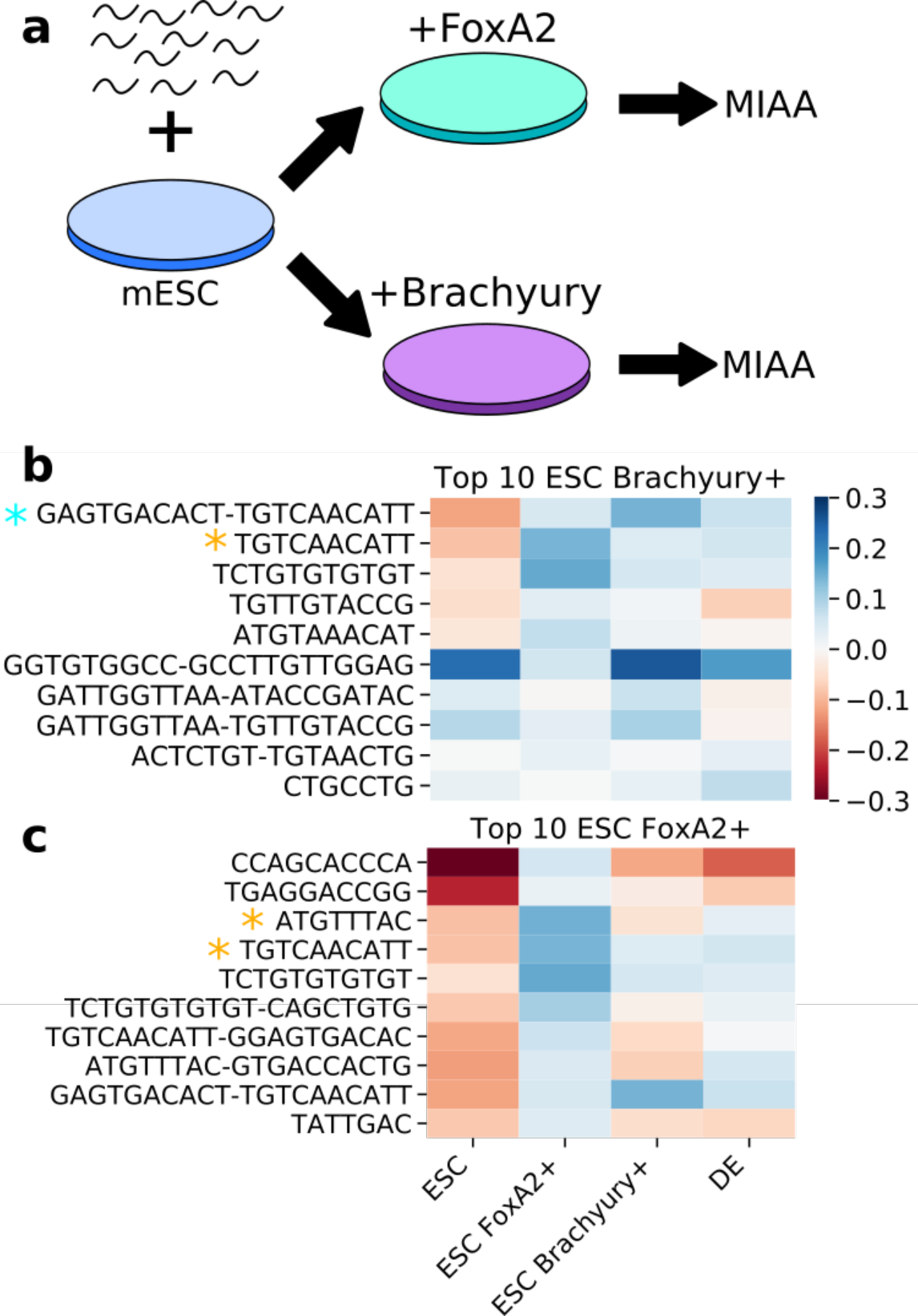
Overexpression of definitive endoderm pioneer transcription factors results in changes to certain motifs representing DNA binding. A) Synthetic phrase library is integrated into stem cells and FoxA2 and Brachyury are over-expressed. B) Heatmap of regression weights of top motifs and motif pairs which increase accessibility in Brachyury compared to stem cells. Blue star indicates motif visually matches Bracyury homodimer in -/+ orientation which is enriched in ChIP-seq peaks. Yellow star indicates motif is statistically significantly enriched in ChIP-seq peaks of Brachyury binding in mouse definitive endoderm cells (p< 0.05 HOMER motif enrichment with Benjamini-Hochberg correction). C) Heatmap of regression weights of top motifs and motif pairs which increase accessibility in FoxA2 compared to stem cells. Star indicates motif is statistically significantly enriched in ChIP-seq peaks of FoxA2 binding in mouse definitive endoderm cells (p< 0.05 HOMER motif enrichment with Benjamini-Hochberg correction).

Similarly, we examined the motifs with the greatest increase in accessibility upon FoxA2 over-expression and found that the third and fourth top motifs were enriched in sequences from FoxA2 ChIP-seq peaks (p < 0.05 under Benjamini-Hochberg multiple hypothesis correction; Figure 4c). FoxA2 over-expression results in more changes from stem cell motif accessibility profiles (Supplementary Figure 10), which is consistent with FoxA2 over-expression resulting in more changes to gene expression (Supplementary Figure 11). The TGTCAACATT motif was enriched in both Brachyury and FoxA2 ChIP-seq, which is likely a result of a common “AACA” subsequence in both FoxA2 and Brachyury binding motifs. We also found that both FoxA2 and Brachyury over-expression resulted in changes in the effects of motifs that generally brought cells closer to the MIAA profile of definitive endoderm (Supplementary Figure 9). Thus, differential accessibility as measured by MIAA accurately matches expected changes in chromatin accessibility induced by transcription factor over-expression.

### IV Exploration of ordering of pioneer transcription factors uncovers subtle TF-TF interactions

Finally, we explored if any transcription factor interactions influencing accessibility were the result of specific TF-TF interactions beyond additive effects. We first trained four linear regression models on each cell type using the phrases from our motif library with models using either 1) GC-content alone, 2) an additive mode with linear effects for each motif, 3) a log-additive model with multiplicative effects for each motif, or 4) a conditional model for which each motif or motif pair has its own effect. Of the models which include motif effects, an additive or a log additive model are the same complexity in terms of number of parameters (one for each motif) with a different assumption in whether we expect the effect of each transcription factor to be added or multiplied when they are in the same phrase. A conditional model is more complex, and expects that each transcription factor pairing can have a unique impact on accessibility. We found that a conditional model best explained the variance of held-out test data over 8-fold cross validation (p < 0.01 by one-tailed Wilcoxon Signed-Rank Test); Supplementary Figure 12).

However, the design of our original library and the close proximity between motifs meant it was possible that pairs of motifs represent a substrate for new proteins to bind and therefore favor a non-linear model. To probe non-linear interaction effects over a constrained set of known factors, we designed a new library from the consensus binding motifs of stem cell pioneer transcription factors Oct4, Sox2, and Klf4 (Figure 5a) and the definitive endoderm transcription factors FoxA2, Sox17, and Gata4 (Figure 5d). We note that the consensus motifs for Sox17 and Sox2 are highly similar, sharing a common sequence (CATTGTTT), so it is highly likely that both Sox factors and possibly others bind to both motifs tested. We tested homotypic phrases with 1, 2, or 3 instances of a motif and heterotypic phrases with combinations of motifs with every possible ordering of the motifs. We found that single motif instances were able to significantly increase accessibility as compared to shuffled phrases for 2/6 transcription factors (Sox17 and Gata4) but were rarely able to make DNA significantly differentially accessible (Supplementary Figure 13). In our phrases containing 2 motif instances, 17/18 significantly increased accessibility as compared to shuffled phrases in at least one cell-type (Supplementary Figure 14), indicating that MIAA is capable of reliably detecting accessibility changes resulting from a minimum of two motif instances and that all of the six motifs utilized are sufficient to open chromatin in at least one cell type. We then tested for differential accessibility under a distance of 6nt and 20nt between motifs, which we selected based on literature supporting preferential distances between Sox2 and Oct4 and between Klf4 and Oct4^46^, and we found that none were significantly different under multiple hypothesis testing. Though we chose canonical motifs for factors well-known in the literature to be associated with stem cells and endoderm, these motifs themselves are insufficient to drive differential accessibility, instead driving accessibility indiscriminately in both cell types. There are several possible explanations for this result. First, certain factors such as Oct4 are expressed in endoderm as well as in stem cells. Second, certain motifs may be shared by multiple members of a transcription factor family. In fact, it has been shown that FoxD3 binds in stem cells to motifs that will eventually become occupied by Foxa2 in endoderm^47^. This same effect likely holds for Sox2 and Sox17 as well, among others, given the similarity of their motifs.

**Figure 5.**
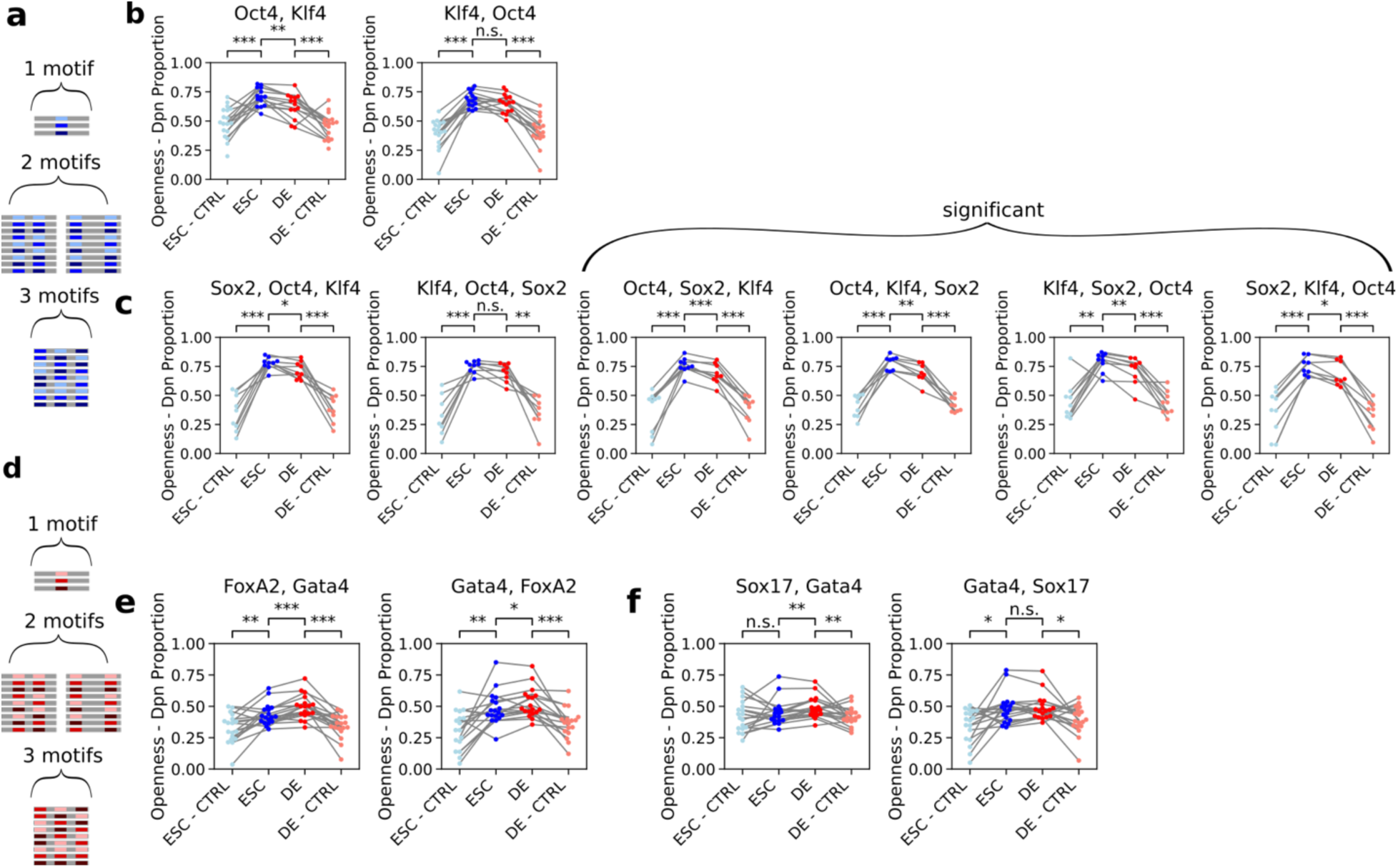
Pioneer motifs have preferential spatial ordering. All significance values reported from paired t-test. A) Phrase construction from stem cell pioneer transcription factors Oct4, Sox2 and Klf4. B) Oct4 and Klf4 motifs have preferential ordering resulting in differential accessibility. C) Preferential ordering of phrase conformations containing all three stem cell pioneers for differential accessibility. D) Phrase construction from stem cell pioneer transcription factors Gata4, Sox17 and FoxA2. E) Preferential ordering of FoxA2 and Gata4 results in differential accessibility. F) Preferential ordering of Sox17 and Gata4 results in differential accessibility.

To determine if particular combinations and/or orientations could drive differential accessibility, we then tested all conformations with 1, 2, or 3 motif instances for induction of accessibility and differential accessibility. Overall, we found that 35/42 conformations significantly increased accessibility as compared to shuffled versions in at least one cell-type, and 15 out of 42 motif conformations were statistically significant for differential accessibility induction after multiple hypothesis correction. Of these 15 conformations inducing differential accessibility, 9 are conformations of 2 motifs and 6 are conformations of 3 motifs, and 10 out of the 15 differentially accessible motif conformations are heterotypic (Supplementary Figure 15). We note that some of the bias toward 2 motif phrases could be due to the fact that 2 motifs phrases have more samples as they were tested over two possible distances, compared to 1 or 3 motif instance phrases.

For stem cell factor binding motifs, we found that one ordering of Oct4 and Klf4 differentially opens chromatin, while the other opens chromatin equivalently in both cell types (Figure 5b).We also found 4 out of 6 phrases that contained all three stem cell pioneer motifs were differentially accessible, and that the order of these motifs had an impact on the level of differential accessibility (Figure 5c) that was somewhat consistent with effects of order of these motifs on genomic accessibility as measured by stem cell DNase-seq (Supplementary Figure 16; see Supplementary Methods for details).

Among endoderm factor motif combinations, we found that particular orientations of FoxA2 and Gata4 (Figure 5e) and of Sox17 and Gata4 (Figure 5f) promoted more differential accessibility that again were consistent with the effects seen from endoderm DNase-seq (Supplementary Figure 16; see Supplementary Methods for details). Previous studies have implicated Gata family members as pioneering transcription factors^1^, including evidence that Gata4 interacts with FoxA2 to drive accessibility changes during endoderm differentiation^30^.

Finally, we investigated whether any of these motif combinations showed evidence of non-linear effects on accessibility by evaluating GC-content only, additive effect, multiplicative effect, and conditional interaction effect regression models with 25-fold cross validation on their ability to predict the stem cell and definitive endoderm MIAA-measured accessibility (See Supplementary Methods for details). Evaluating over all test phrases, stem cell accessibility and definitive endoderm accessibility appears to be equally well fit by either an additive or a multiplicative model (Supplementary Figure 17). Introducing conditional TF-TF interaction effects for each possible conformation of transcription factors did not significantly increase overall prediction power. We then evaluated individual transcription factor combinations for a significant improvement in model fit for stem cell accessibility prediction, definitive endoderm accessibility prediction, or differential accessibility prediction (Supplementary Table 2). At the level of individual combinations, most are best fit by an additive model (26/30 stem cell, 26/30 endoderm). While this analysis did not recover any strongly conditional or multiplicative interactions, the subtle effects of motif order on differential chromatin accessibility suggest that MIAA has the capacity to measure such interaction effects. In sum, the spatial order of transcription factors can influence chromatin accessibility, albeit in subtle ways. MIAA is well-suited to identifying such subtle effects through its ability to test many permutations of transcription factor combinations in a controlled sequence context. Additionally, we find evidence that chromatin accessibility in stem cell and definitive endoderm obey a mostly additive logic with respect to the inputs of individual transcription factor motifs. Such findings are useful in guiding computational models of cell type-specific chromatin accessibility.

## Discussion

The Multiplexed Integrated Accessibility Assay (MIAA) is a new assay for measuring changes in chromatin accessibility caused by short DNA phrases integrated into a fixed locus in the genome. Most prior approaches to understanding the control of chromatin accessibility have used correlative approaches that identify genomic DNA sequences that tend to coincide with accessible chromatin in a particular cell type^9,28,48–50^. Several studies have screened for natively occurring SNPs in cohorts of patient cell lines, identifying “DNase-QTLs” for which the single nucleotide change induces a change in chromatin accessibility^51,52^, revealing motifs whose disruption is enriched in such variants. MIAA enables screening of an arbitrarily large and diverse library of sequences for their impact on chromatin accessibility.

We applied MIAA to understanding the effects of motifs on differential accessibility between stem cell and definitive endoderm cell states using a number of distinct experimental designs. Through the use of native genomic 100-nt phrases transplanted to a fixed locus, we identify an intriguing distinction between how a set of natively stem cell-specific and endoderm-specific sequences achieved differential accessibility. The natively endoderm-accessible sequences opened chromatin more in endoderm than in stem cells and more than their shuffled versions on average, suggesting the presence of binding sites for endoderm-specific accessibility-promoting transcription factors. On an individual level, only a subset of sequences act in this way, suggesting that 100-nt is only sometimes sufficient to recapitulate the chromatin accessibility status of native regulatory elements and offering an assay to determine which native genomic regulatory sequences are sufficient to encode differential accessibility.

Intriguingly, we found a distinct pattern in the natively stem cell-accessible sequences. In this cohort of sequences, MIAA accessibility was higher in stem cells than in endoderm as expected; however, there was no difference between the stem cell accessibility between the sequences and their shuffled counterparts. Instead, the accessibility in endoderm was reduced as compared to shuffled controls, suggesting that differential accessibility of these sequences was primarily achieved through binding sites that depress accessibility in endoderm. We found suggestive evidence that E-box binding sites used by epithelial-mesenchymal transition driver TFs such as ZEB factors may be responsible for this repression. Because the native genomic sites were selected based on predicted optimal differential accessibility from a sequence-based model using DNase-seq regions, it is striking to have detected such a consistent difference in the mechanism of achieving differential accessibility, and it will be intriguing to explore a larger cohort of cell type-specific sequences to determine which mechanism is more common.

In order to identify causal motifs and transcription factors involved in mediating differential chromatin accessibility, we then focused on exploring phrases containing various combinations of sequence motifs. We show that, independently of binding motifs, higher GC-content increases accessibility. In MIAA, we can confirm this to occur in the absence of transcription factor binding motifs because of our use of shuffled versions of each designed phrase. While it is formally possible that this GC effect is an artifact of the use of Dam methylase, we show that native genomic accessible regions also show elevated GC-content, and it has been reported that transcription factors and DNase hypersensitive regions are also enriched in GC-rich regions^7^.

In spite of its importance, predicting MIAA chromatin accessibility of held-out phrases purely based on GC-content yields poor results, while much better results are achieved by accounting for binding motifs. Of the motifs that can be confidently matched to known transcription factor families, our results are consistent with the action of known tissue-specific pioneer factors including Sox2 and Oct4 in stem cells and FoxA2 and Brachyury in endoderm^8,53^. We are able to widen this list of cell type-specific pioneer TFs, demonstrating the power of MIAA to identify both robust and subtle effects of motifs on accessibility with high statistical confidence through the use of matched shuffled control sequences. We further confirm the role of FoxA2 and Brachyury in endoderm-specific chromatin opening by demonstrating that over-expression of these definitive endoderm transcription factors in stem cells can increase MIAA-measured accessibility significantly in phrases with DNA-binding motifs recognized by these factors. We then designed a library using consensus motifs of several pioneer factors in all possible combinations and orderings, from which we provide evidence that a single binding site is sufficient to open chromatin and as few as two binding sites is sufficient to differentially open chromatin between two cell types. We also found that for motifs known to bind to both stem cell and definitive endoderm transcription factors, motif order has a subtle effect on accessibility, which provides support for specific transcription factor interactions driving accessibility change. This result illustrates the complexity of differential accessibility induction, which cannot simply be distilled to the presence of consensus motifs for differentially expressed transcription factors. In addition to the re-use of genomic motifs by different members of the same transcription factor family in different cell states^47^, certain transcription factors such as those in the Sox and Oct family can exhibit profoundly distinct binding to specific dimeric motifs that differ in subtle ways^54^. MIAA offers an exciting new way to explore subtleties that influence transcription factor binding logic such as motif ordering, spacing, and dimeric motifs in a controlled genomic setting.

We also found that, for the vast majority of motif arrangements (26/30 in stem cells, 26/30 in endoderm), a model in which motif instances have an additive impact on chromatin accessibility best fits the observed MIAA data. While our analysis did not support strong conditional interactions between transcription factors, we observed subtle effects of motif order and strong change in accessibility by a motif pair matching a Brachyury dimer when Brachyury was over-expressed, suggesting that MIAA has the capacity to measure the effects of transcription factor interactions on accessibility. Predicting differential accessibility from DNA sequence has been a much more difficult task^27,55,56^, and one possible reason is that more conditional logic is used. Due to the low number of instances of motif co-occurrence in cell type-specific accessible regions, conclusions about the impact of conditional logic or preferred orientation on accessibility would be difficult for models such as DeepAccess to learn from the genome alone. The ability of MIAA to obtain sensitive measurements of the effects of specific motif combinations on differential accessibility by designing more instances than occur in the genome in a controlled fashion makes MIAA a valuable tool in training accurate predictive models of chromatin accessibility.

There are many directions for future work, including a deeper examination of the impact of genomic integration site on local DNA accessibility as well as a further investigation into features such as motif spacing which are likely to impact transcription factor interaction logic. Another possible application of MIAA is to understand chromatin accessibility during differentiation by taking measurements at multiple timepoints to discover novel transcription factor logic in developmentally relevant cell types.

## Supporting information

MIAA-phrase-counts-library

MIAA-supplementary-methods

MIAA-supplementary-figures

## Contributions

JH designed and implemented DeepAccess and performed analysis of DNase-seq, ChIP-seq, and RNA-seq data. J.H., K.K., D.K.G., and R.I.S. designed MIAA libraries. R.I.S. and B.B. conceived and carried out MIAA experiments. J.H. and K.K. performed processing and computational analysis of MIAA results. J.H., K.K., D.K.G., and R.I.S. contributed to interpretation of results. J.H., D.K.G., and R.I.S. wrote the manuscript with input from all authors.

## Acknowledgments

We thank MIT BioMicro Center and the Harvard Medical School Biopolymers Facility for reagents and technical assistance and members of the Gifford lab, Sherwood lab, and Wichterle lab for helpful discussions. We thank Dr. Hynek Wichterle for comments on the manuscript. We gratefully acknowledge funding from 1RO1HG008363 (D.K.G.), 1R01HG008754 (D.K.G.), 1R01NS109217 (D.K.G.), 1K01DK101684 (R.I.S.), the Human Frontier Science Program, NWO, American Cancer Society, and Qatar Biomedical Research Institute (R.I.S.), and National Science Foundation Graduate Research Fellowship (1122374) (J.H.).

## Methods

### Embryonic stem cell line generation, culture, and endoderm differentiation

All experiments were performed in dual LoxP site-containing and rtTA-expressing 129P2/OlaHsd mouse embryonic stem cells (mESC), which were cultured according to previously published protocols^57,58^. Cells were regularly tested for mycoplasma. Cre/LoxP site-specific integration of a Rarg-DamR126A fusion construct (subcloned from a previously published Tcf7l2-DamN126A mutant construct^35^ was performed followed by G418 (Life Technologies) selection (350 *μ*g/mL) for one week to generate the mESC line used in all experiments. To generate sublines with inducible expression of FoxA2 or Brachyury, full-length mouse cDNAs for these genes were cloned into a Tol2 expression plasmid containing a 7X TRE promoter and Blasticidin resistance^1^. Co-transfection of Tol2 transposase with TRE-Foxa2/Brachyury plasmids was performed followed by selection with 10 ug/mL Blasticidin for one week.

mESCs were maintained on gelatin-coated plates feeder-free in mESC media composed of Knockout DMEM (Life Technologies) supplemented with 15% defined fetal bovine serum (FBS) (HyClone), 0.1mM nonessential amino acids (NEAA) (Life Technologies), Glutamax (GM) (Life Technologies), 0.55mM 2 *β*-mercaptoethanol (Sigma), 1X ESGRO LIF (Millipore), 5 nM GSK-3 inhibitor XV and 500 nM UO126. To induce differentiation, cells were grown for 48 hours in mESC media without GSK3 inhibitor XV and UO126 and then seeded at 10^4 cells/cm2 in mESC media without GSK3 inhibitor XV and UO126. After 24 hours, media was replaced with serum-free differentiation media: Advanced DMEM supplemented with N-2, B27 Supplement without vitamin A, and Glutamax (Thermo Fisher). After 24 hours, media was replaced with differentiation media: Advanced DMEM with 2% ES-tested FBS and Glutamax. To induce endoderm differentiation, cells were treated for 24 hours in differentiation media with 50 ng/mL E. coli-derived Activin A (Peprotech) for 24 hours to produce mesendoderm, then cells were fed with Advanced DMEM with 2% FBS, Glutamax, 50 ng/mL Activin A and 1 µM Dorsomorphin (Sigma) for 48 hours. Induction of Rarg-DamN126A expression was performed by treatment of mESCs or mESC-derived endoderm with 2 ug/mL Doxycycline (Sigma) for 24 hours prior to genomic DNA collection using the Purelink Genomic DNA miniprep kit (Life Technologies).

### Oligonucleotide library integration and NGS library preparation

Oligonucleotide libraries were ordered in the following format from Twist Biosciences: CACTCAGTACTTTGTCCGTGCTGAC [100-nt phrase] AGATCGGAAGAGCGTCGTGTAGGGA

Note that this sequence includes a reverse complement of the Illumina read 1 sequencing primer to facilitate NGS. All PCR amplification was performed using 2X Q5 UltraII Mastermix (New England Biolabs).

Genomic integration was performed by PCR amplification of oligonucleotide libraries with the following homology arm primers:

RARRXR1 locus (primary locus used in this work)

RARRXR1_library_fw

CCTGGTCCAGACACTCATTCTCAAGCTTCCTCATGCTCTTGTGGGAAGCATAGATGCTTTCAGAG CA CTCAGTACTTTGTCCGTGCTGAC

RARRXR1_library_rv

CTGTGAGGCTGGTGGAAGACCACAAACAGGGGAGGGTCATGGAGAGGTCAGGGGTTGCCAACAAAGC TCCCT ACACGACGCTCTTCCGAT

uCd8 locus (secondary locus used in this work)

uCd8_library_fw

CGAATCACTCCATGTGAGTATCACAGAACGGGTGCAGGAGATCAGTTGCTGT GATGGATAGACACAG TCCCTACACGACGCTCTTCCGAT

uCd8_library_rv

ATCCGCCCTGAAGCAGGCAGCAGAGCAGATGCTCTGAGATGCTTGCTTTCTGTAGCCCAGGTGTG CACTCAGTACTTTGTCCGTGCTGAC

for 35 cycles, using 60 second 72 deg extension in each cycle to reduce GC bias. For each 15 cm plate to be electroporated, 1% of the library was amplified in 500 uL PCR volume. MinElute purification with 250 uL PCR product/column was performed, eluting in 10 uL/column and pooling the two columns’ worth of product for each plate to be electroporated.

*Sp*Cas9 gRNAs were cloned into a Tol2-transposon-containing gRNA expression plasmid (Addgene 71485) ^59^.) using BbsI plasmid digest and Gibson Assembly (New England Biolabs). Spacer sequences were as follows:

RARRXR1: GAGCAGGTGACAATTTCAGA

uCd8: GTAGCCCAGGTGTGCAGGCT

For each 15-cm plate to be electroporated, we used 40 ug p2T CBh Cas9 BlastR (Addgene 71489), 40 ug gRNA plasmid, and minElute-purified product from 500 uL PCR.

Electroporations were performed in 2-4 biological replicates into p2L RAR-DamA126 cells by mixing the listed DNA amounts with 5-20*10^6 cells in 300 ul EmbryoMax Electroporation Buffer (Millipore) and electroporating in a 0.4-cm electroporation cuvette using a BioRad electroporator at 230 V, 0.500 mF, and maximum resistance. Electroporated cells were plated in mESC media supplemented with 7.5 mM Y-27632 (Tocris). From 24 to 72 hr after electroporation, media was refreshed daily with mESC media supplemented with 10 ug/ml Blasticidin (Life Technologies) and 66 ug/ml (1:666) Hygromycin (Cellgro). Cells were grown for 5-8 days after electroporation to obtain adequate quantities for Doxycycline treatment and endoderm differentiation.

Genomic DNA library preparation was performed by performing DpnI and DpnII digests using the following conditions:

DpnI digest: 10 ug genomic DNA + 10 uL CutSmart Buffer (New England Biolabs) + 2 uL DpnI (New England Biolabs) + up to 100 uL water.

DpnII digest:20 ug genomic DNA + 20 uL DpnII buffer (New England Biolabs) + 4 uL DpnII 9New England Biolabs) + up to 200 uL water. Reactions were incubated for 16-24 hours at 37 deg then 80 deg for 30 min. Digested genomic DNA product was used directly as input to library prep.

Library prep was performed by 3 successive PCRs. In the first PCR, the entire DpnI-digested genomic DNA was used in a 400 uL PCR rxn, and the entire DpnII-digested genomic DNA was used in a 800 uL PCR rxn. 13 cycles of PCR1 (Q5 UltraII mastermix, Ta=66, 60 second 72 deg extension per cycle) was performed using the following primers:

RARRXR1 locus:

Library_PCR1_fw TCAGTACTTTGTCCGTGCTGAC

RARRXR1_PCR1_rv (for RARRXR1) GTCCACCCTTCCTGTCTGTAC

uCd8 locus:

uCd8_PCR1_fw CCGGTGGGGTCTCAGTGTTAACC

Library_PCR1_rv TCCCTACACGACGCTCTTCCGAT

PCR 1 was purified using PCR purification in a single column, eluting in 50 uL. PCR2 was performed for 7 cycles (Q5 UltraII mastermix, Ta=66, 60 second 72 deg extension per cycle) using 22.5 uL of PCR1 product in a 100 uL reaction with the following primers:

Library_Read1stub ACACTCTTTCCCTACACGACGCTCTTCC

Library_Read2stub GTTCAGACGTGTGCTCTTCCGATCTCAGTACTTTGTCCGTGCTGAC

PCR 2 was purified in a single column, eluting in 50 uL.

qPCR (Q5 UltraII mastermix with EvaGreen (Biotium), Ta=66, 60 second 72 deg extension per cycle) was used to determine PCR3 cycle counts using the following primers:

Read1_noindex

AATGATACGGCGACCACCGAGATCTACACTCTTTCCCTACACGACGCTCTTCCGATC

T

Read2_noindex

CAAGCAGAAGACGGCATACGAGATGTGACTGGAGTTCAGACGTGTGCTCTTCCGA

TCT

PCR3 was performed by subtracting 2-3 cycles from the Ct and using 2.5 uL of a mix of distinct indexed primers from the NEBNext Dual Index Kit for Illumina (New England Biolabs) using 22.5 uL PCR2 product in a 50 uL reaction volume (Q5 UltraII mastermix with EvaGreen (Biotium), Ta=66, 60 second 72 deg extension per cycle). PCR3 products were PCR-purified and Tapestation (Agilent) was used to quantify and pool samples for NGS.

Illumina Nextseq was used using the following read lengths:

Read 1: 150 nt

Index 1: 8 nt

Index 2: 8 nt

### Phrase library design

We ran a pilot experiment and from the pilot identified 6 native genomic sequences of size 100nt, that did not drive differential accessibility with MIAA but varied in GC-content. We randomly perturbed these native sequences 3 times each to obtain a total of 24 neutral sequence backgrounds. For our first experiment, we took each background and inserted either 1 motif 7 times (positions 2, 16, 30, 44, 58, 72, 86) or 2 motifs where motif 1 is inserted 4 times (positions 2, 30, 58, 86) and motif 2 is inserted 3 times (positions 16, 44, 72). For our second experiment, we limited ourselves to 9 backgrounds which we expected to have high reproducibility to the set of 24. In this experiment we test sequences of size 70nt. Using the consensus sequences of known ES pioneers (Oct4, Sox2, Klf4) or DE pioneers (FoxA2, Gata4, Sox17), we inserted 1, 2, or 3 motifs into each sequence. We tested homotypic phrases consisting of one unique motif, as well as heterotypic phrases enumerating all possible motif orders.

### Phrase library sequencing and processing

Reads were mapped to library phrases by taking the reverse complement to the raw read, where the first N nucleotides (between 70 and 100 based on the size of the designed phrase) are the designed variable phrase. Perfect matches were counted using a custom R script. Reads were normalized to reads per million over the total number of reads in the digest. Phrases were kept if they had a threshold number of total normalized reads over all replicates, based on the observation of high standard deviation at low total read counts. Threshold was selected based on visual inspection (Supplementary Figure 18). Once reads were normalized and high variability phrases filtered, Dpn proportions were computed as DpnII/(DpnI + DpnII) to represent relative openness of the phrase.

### DeepAccess Model and Motif Importance

We obtain DNase-seq regions using the 100nt centered at the MACS2 narrow peak call. Accessibility prediction is treated as a multi-task classification problem, where each genomic sequence (100bp) is associated with a two-dimensional bit vector representing whether the sequence is open in each cell type (ESC and DE). We trained an ensemble of 10 convolutional neural networks. To limit training difficulty, the convolutional filters of the first hidden layer were set to the position weight matrices of the 664 HOMER^49^ motifs of known transcription factors. In the ensemble, one convolutional neural network no hidden layers, followed by a global max-pooling layer, representing a logistic regression classifier with HOMER^49^ motif features. The other 9 convolutional neural networks have a second convolutional layer followed by a global max-pooling layer which allows them to integrate combinatorial and spatial relationships between HOMER^49^ motifs. The fully-connected output layer present in all neural network architectures contains two neurons with a sigmoid activation function that returns a value between 0 and 1 for each neuronal activation, which represents the probability of the predicted DNA “openness” in each of the two cell-types. DeepAccess is trained on a balanced dataset with 400,000 sequences across four possible classification scenarios of a sequence 1) open in endoderm cells and closed in stem cells, 2) open in stem cells and closed in endoderm cells, 3) open in both cell types, 4) closed in both cell types. A test set of 22,357 sequences is held out for performance evaluation.

We extracted motifs from DeepAccess by applying gradient ascent to score each neural network feature input by its importance for predicting the output^60^ and multiply times the input (a one-hot encoding of the DNA sequence) since gradients will assign non-zero values to DNA characters not present in the sequence. In order to obtain sequence importance for features that drive accessibility differentially between definitive endoderm and stem cells, we set the gradient loss to the difference between the predicted openness of two cell types. We also used a smoothed form of gradient ascent, which averages the gradient over multiple noisy inputs^61^. We took the weighted average importance over the ensemble of neural networks based on performance accuracy on held-out test data. We then selected windows of size 10 with the highest average saliency over a set of 5,000 training sequences and used those as the DeepAccess-derived motifs. We also extracted the top motifs with the highest increase in saliency of differential accessibility between the CNN without trainable hidden layers and the CNNs with hidden layers, which represents motifs that gain importance from the CNNs that learn relationships between motifs.

## Code availability

Code for DeepAccess accessibility prediction and motif extraction is available at https://github.com/gifford-lab/DeepAccess. Code for MIAA library processing and producing all manuscript figures is available at https://github.com/gifford-lab/MIAA-analysis.

## Data availability

All previously published data analyzed for this study are listed in Supplementary Table 3. The data generated for this study have been deposited in NCBI’s Gene Expression Omnibus and are accessible through GEO Series accession number GSE145920. Raw MIAA read counts are available as Supplementary Data.

