## Supplementary material for "Identification of determinants of differential chromatin accessibility through a massively parallel genome-integrated reporter assay": MIAA-supplementary-methods

**DNase-seq processing**

DNase-seq reads were aligned to mm10 with bwa aln with the parameters -n 2 -k 2 -t 4. Bams were sorted and paired reads processed with samtools sort and samtools fixmate. Peak accessible regions and signal tracks were generated from aligned reads using the Kundaje lab pipeline (<https://github.com/kundajelab/atac_dnase_pipelines>) using MACS2 peak calling. For each cell type, we define DNase-seq accessible regions as 100nt regions centered at the MACS2 narrow peak calls from the pooled replicate data. Cell type-specific accessible regions are those which do not physically overlap the other cell type.

**RNA-seq processing**

RNA-seq data was downloaded as fastqs. Adaptors were trimmed with TrimGalore version 0.6.2. Trimmed reads were aligned to mm10 and gene read counts were computed with RSEM^1^ version 1.3.0. Differential expression analysis was computed using rsem-run-ebseq and rsem-control-fdr with an fdr threshold of 0.05.

**ChIP-seq processing**

ChIP-seq reads were aligned using bowtie2 version 2.2.9 with the parameters –fast. Duplicate reads were removed with samtools rmdup. Binding events were called with GPS version 3.4 with parameters --d $gemdir/Read_Distribution_default.txt --f SAM. Motif analysis was performed using HOMER^2^ findMotifsGenome.pl with parameters -bg brachyury_chip-union.downstream.bed -size 50 -h. Where brachyury_chip-union.downstream.bed is a custom background bedfile of regions immediately downstream of the GPS ChIP binding region. K-mers from phrase library design were tested by first converting k-mers to motifs using HOMER’s^2^ seq2profile.pl, allowing for up to 2 base mismatches.

**Brachyury Dimer analysis**

See Supplementary methods ChIP-seq processing for methods for ChIP binding event calling and motif enrichment. After motif enrichment, the Brachyury binding sites were scanned for the top Brachyury motif using HOMER^2^ findMotifsGenome.pl with the parameters -size 50. HOMER^2^ output was processed to identify Brachyury binding regions that consisted of 2 motif instances.

**KEGG Pathway Analysis of Zeb2 sites**

Zeb2 motif matches were computed using HOMER^2^ findMotifsGenome.pl on 100nt regions open in stem cells and not definitive endoderm. KEGG enrichment was performed on all Zeb2 motif sites in ESC regions using HOMER^2^ annotatePeaks.pl with the parameters -go $OUTPUT_GO_DIR and otherwise default parameters.

**Statistical test for differential accessibility of k-mers**

For each hypothesis (k-mer or combination of k-mers) a paired t-test and Wilcoxon signed-rank test was computed between the accessibility of phrases in stem cell and accessibility of phrases in definitive endoderm cells. The p-values for each test were adjusted for multiple hypothesis testing by Benjamini-Hochberg procedure with error rate < 0.05. For all phrases which were significant under both t-test and signed rank, a second statistical test was computed within each cell type comparing phrases to paired shuffled controls using paired t-test and Wilcoxon signed-rank test. The p-values for the two cell types were compared, and the lower p-value selected. These tests were adjusted for multiple hypothesis testing by Benjamini-Hochberg procedure with error rate < 0.05, and any hypothesis which passed significance was then considered differentially accessible because it passed both differential between the two cell types as well as differential compared to shuffled controls.

**KMAC motif discovery**

To run KMAC, we take the broad pooled peaks as output from MACS2, and select the top 10,000 which are unique (not physically overlapping) in each cell type. We take the top 10,000 shared peaks as our negative set of sequences. We then run KMAC with the parameters --k_win 100 --k_min 4 --k_max 13 --t 1 --k_seqs 10000 --k_top 10 --gap 4. We selected k-mers from KMAC which showed promise in a pilot MIAA experiment or interesting combinatorial results *in silico* from predictions by our ensemble of convolutional neural networks, DeepAccess.

**Native phrase motif discovery**

We ran HOMER^2^ findMotifs.pl with phrases in cluster passed as the positive sequence set, and the shuffled control library phrases for the background sequence set.

**Phrase k-mer motif discovery**

To obtain motifs from designed k-mers based on MIAA-measured accessibility in stem cells and definitive endoderm, we first ran agglomerative clustering on the regression weights from the k-mers and k-mer pairs. We then extracted all phrases from a given cluster and ran HOMER^2^ findMotifs.pl with phrases in cluster passed as the positive sequence set, and the same number of randomly selected phrases excluding sequences in the cluster as the background sequence set.

**Linear Regression Models**

For visualizing motif accessibility effects, we train linear regression models with the following design matrix, where *r* indicates a replicate-specific offset, *ct* indicates the cell-type indicator, *k* indicates the k-mer or pair of k-mers in the 6k library, and *gc* indicates the GC content of the phrase. Each phrase will have up to 3 non-zero features and no features should be correlated so we apply non-regularized OLS regression.

$$y=\sum_{r} x_{r}\beta_{r}+\sum_{ct} \sum_{k} x_{ct,k}\beta_{ct,k}+{\sum_{ct} \beta_{ct,gc}x}_{ct,gc}$$

For comparing GC-content alone, additive, multiplicative, or conditional regression models, we trained regression models independently for each cell type on the average accessibility over all replicates with the following design matrices:

1) GC-content:

$$y_{avg}=x_{gc}\beta_{gc}$$

2) Additive:

$$y_{avg}=x_{gc}\beta_{gc}+\sum_{k} x_{k}\beta_{k}$$

3) Multiplicative (log-additive):

$$log(y_{avg})=x_{gc}\beta_{gc}+\sum_{k} x_{k}\beta_{k}$$

4) Conditional:

$$y_{avg}=x_{gc}\beta_{gc}+\sum_{k_{pair}} x_{k_{pair}}\beta_{k_{pair}}+\sum_{k_{single}} x_{k_{single}}\beta_{k_{single}}$$

For these models, we evaluated them using k-fold cross-validation by randomly selecting a set of 3 out of 9 (pioneer 2,000 phrase library) or 8 out of 24 (DeepAccess k-mer 6,000 phrase library) backgrounds to hold out as test data.

**Best model fit of transcription factor interactions**

To assess model fit of specific conformations of phrases containing more than one transcription factor instance, we evaluated the mean-squared error of regression models (see Methods – Linear Regression models for details) on only phrases with the specific set of motifs, with 25-fold cross validation, with each cross holding out 3 out of 9 backgrounds as test data. We tested for significant model improvement with one-tailed Wilcoxon Signed Rank Test, in order of complexity (additive, multiplicative, conditional) such that we favor additive or log-additive models over complex models. For example, we first would compare performance of additive model to multiplicative model on phrases containing foxa2 in motif slot 1 and gata4 in motif slot 2. If the multiplicative model did not significantly (p < 0.05) outperform the additive model, the next test was comparing the conditional model to the additive model. If the multiplicative model did significantly outperform the additive model, the next test was comparing the conditional model to the multiplicative model.

**Genomic visualization of DNase-seq**

This visualization was used to compare relative accessibility of regions with different orderings of pioneer motifs in the genome. DNase-seq bigwig signal tracks and peak regions were generated according to Methods - DNase-seq processing. We then ran HOMER^2^ findMotifsGenome.pl on 100nt regions that were called is open in ES cells (for Oct4, Sox2, and Klf4) or DE cells (for FoxA2 and Gata4 or Sox17 and Gata4) using the HOMER^2^ motifs for the respective pioneer TFs. We then select all instances that have the same TF order, and report the mean and 95% confidence intervals of DNase-seq signal.
