## Supplementary material for "Identification of determinants of differential chromatin accessibility through a massively parallel genome-integrated reporter assay": MIAA-supplementary-figures

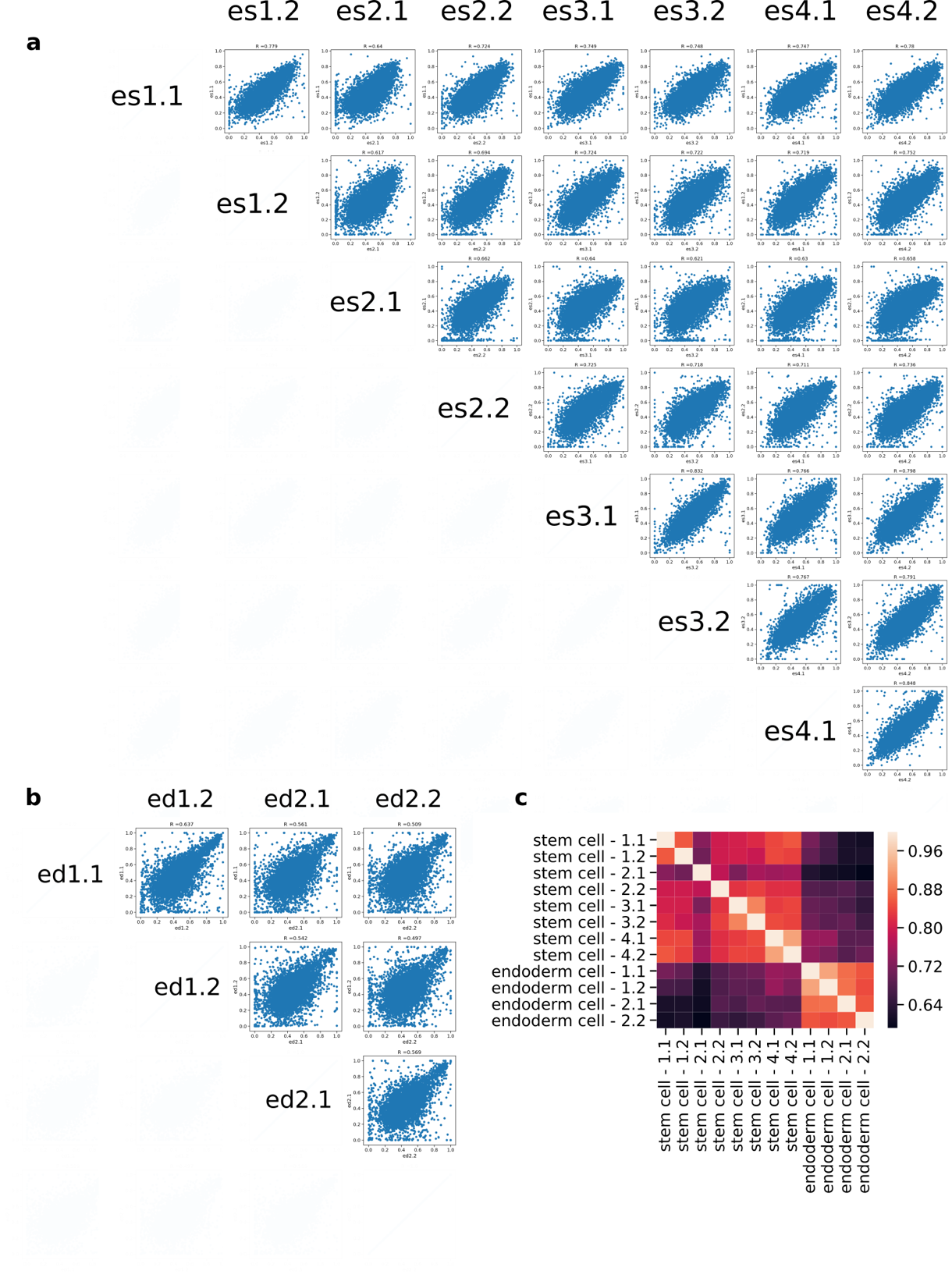

Figure S1. MIAA experiments are highly reproducible. A) Dpn proportion of all stem cell replicate experiments over 6K phrase library. R reported is Pearson correlation coefficient. B) Dpn proportion of all endoderm replicate experiments over 6K phrase library. R reported is Pearson correlation coefficient. C) Heatmap of Pearson correlation of Dpn proportion over universally opening, universally closing and background phrases in both cell types. ­­­

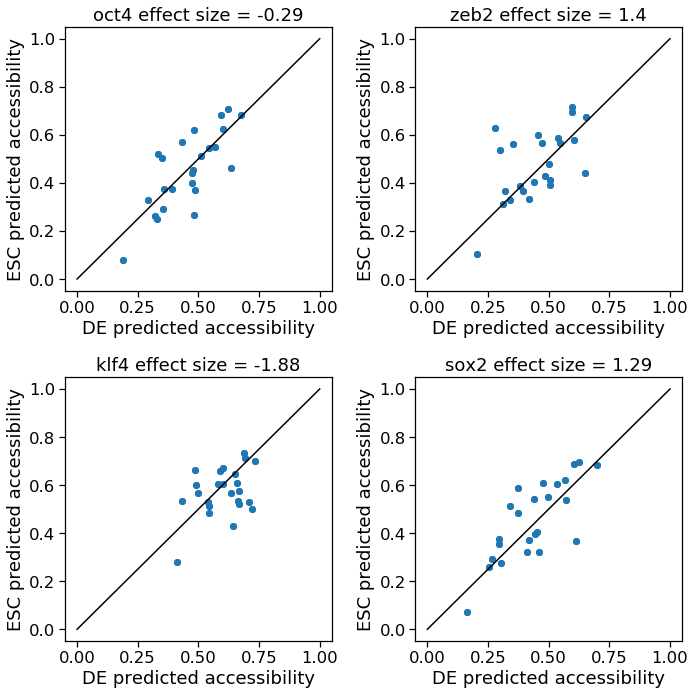

Figure S2. DeepAccess predictions for phrases containing a single consensus sequence for DNA binding motifs on stem cell (ESC) and definitive endoderm (DE) accessibility. Strongest predicted impact on differential accessibility is from Zeb2 (E-box) motif.

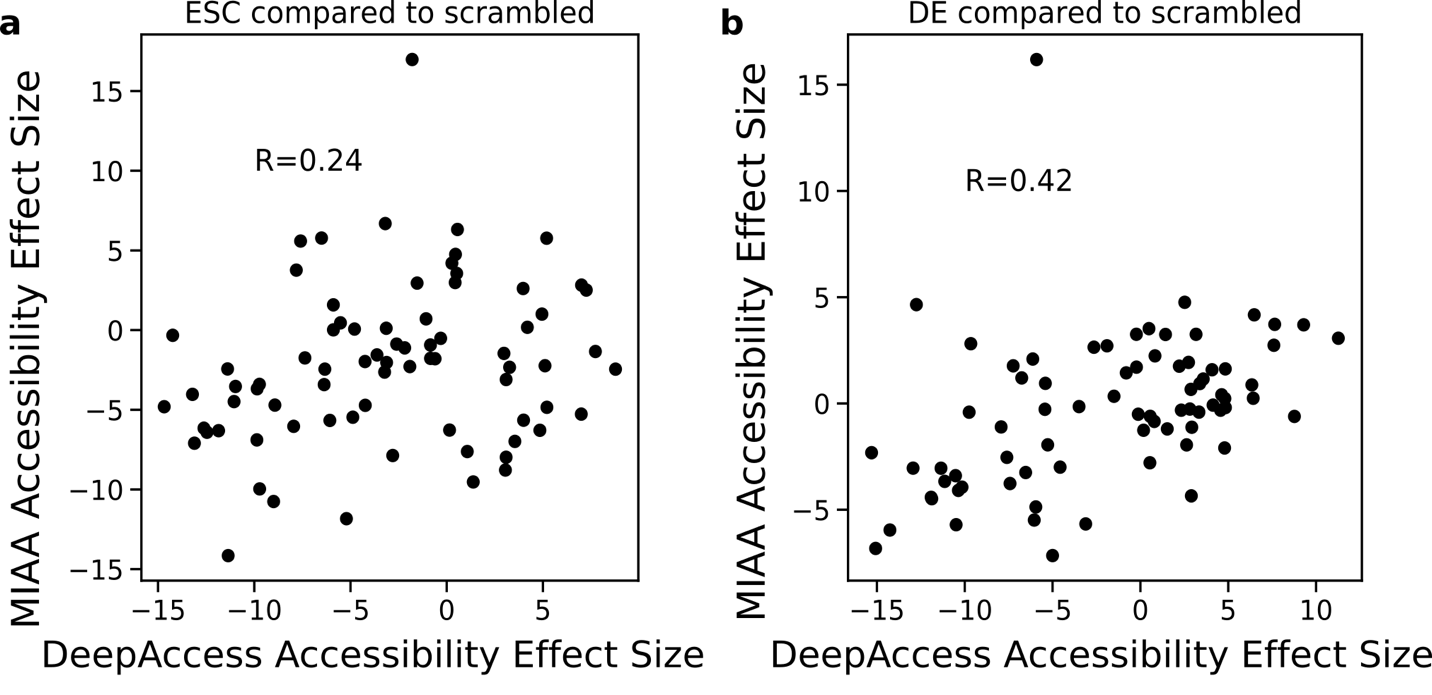

Figure S3. DeepAccess predictions of motif effects from homotypic and heterotypic phrases compared to shuffled controls. Each dot is the measured effect size over 24 backgrounds containing either 6 instances of one motif (homotypic) or 3 instances of two motifs (heterotypic). A) Effects of motifs relative to paired randomly shuffled control phrases in mouse embryonic stem cells in MIAA assay compared to predictions from DeepAccess. B) Effects of motifs relative to paired randomly shuffled control phrases in definitive endoderm cells in MIAA assay compared to predictions from DeepAccess. The correlation reported is the Pearson correlation coefficient (r).

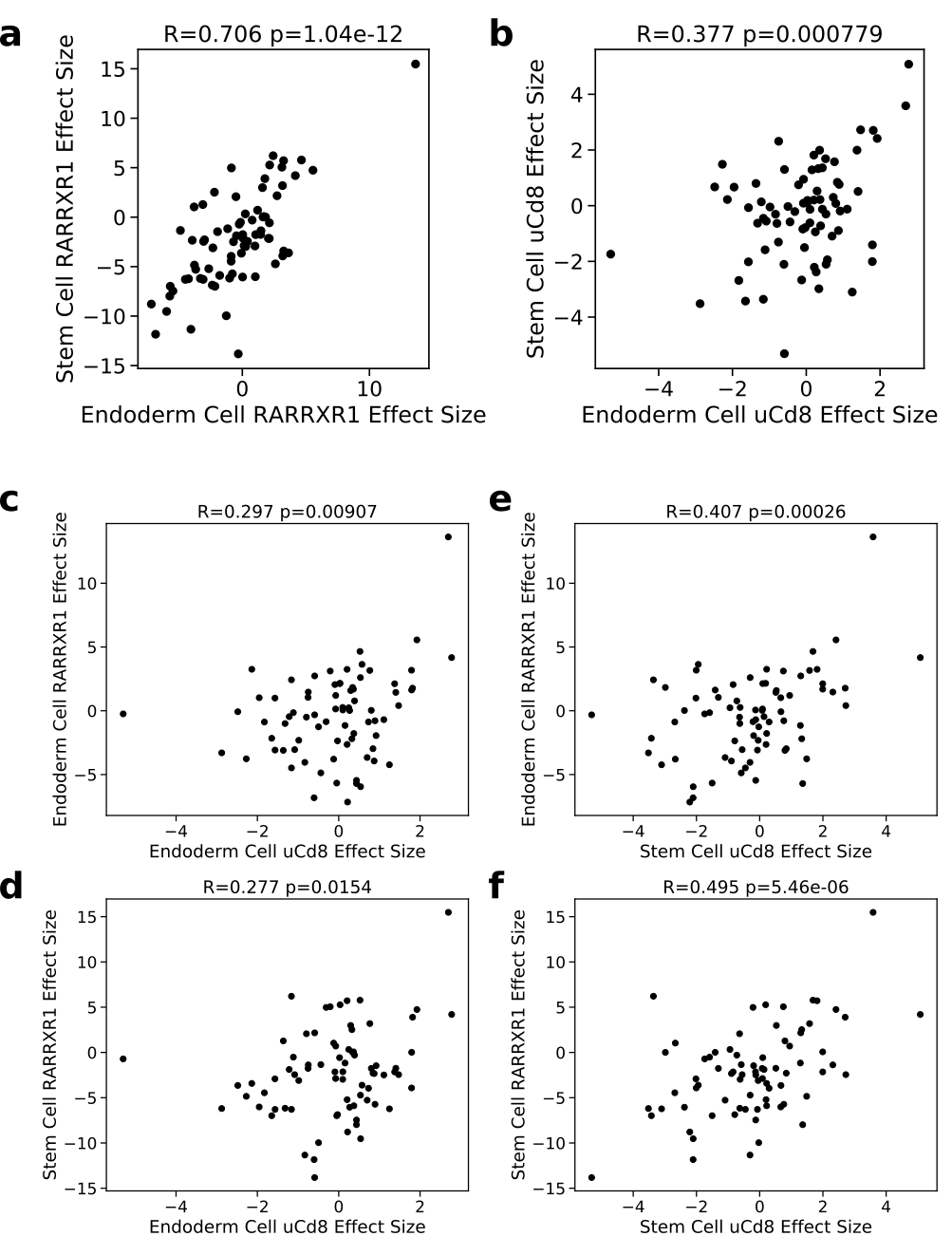

Figure S4. MIAA experiments are reproducible in uCd8 genomic integration locus. Each dot represents the effect size by paired t-test for a motif in 24 neutral backgrounds compared to paired shuffled controls. A) Comparison of effect size between cell types in RARXR1 locus shows high correlation. B) Comparison of effect size between cell types in uCd8 locus shows lower correlation, indicating lower bound on significance of the results. C) Comparison between effect size in definitive endoderm in both loci is higher than between stem cell RARRXR1 and endoderm uCd8 (D), but lower than correlation between endoderm RARRXR1 and stem cell uCd8 (E). Effect size between stem cell RARRXR1 and stem cell uCd8 is highest out of all cross-loci comparisons.

| **kmer** | **model** | **celltype-differential** | **deepaccess-predicted** |
| --- | --- | --- | --- |
| GATTGGTTAA-TGTTGTACCG | DeepAccess | True | True |
| CGCGGCGGCC | DeepAccess | False | N/A |
| GGTGTGGCC-GCCTTGTTGGAG | KMAC | True | True |
| GTCACCTGAG | DeepAccess | True | True |
| ATACCGATAC-GCGCACACCT | DeepAccess | True | True |
| GGTGTGGCC-CTGCCTG | KMAC | True | True |
| TGGAGTGACA | DeepAccess | True | True |
| ATGTTTAC-CGCGGCGGCC | DeepAccess | True | False |
| TGTTGTACCG | DeepAccess | False | N/A |
| GATTGGTTAA-GCGCACACCT | DeepAccess | True | True |
| TGATGTCACT-TGGAGTGACA | DeepAccess | True | True |
| GGCCATTGT | KMAC | True | True |
| GATTGGTTAA | DeepAccess | True | False |
| ACGAATTATC-GTCACCTGAG | DeepAccess | False | N/A |
| CACCAGGGG | KMAC | False | N/A |
| TGCAGTGACA-TGGAGTGACA | DeepAccess | True | True |
| TGTCAACATT-GGAGTGACAC | DeepAccess | True | True |
| GAGTGACACT-TGTCAACATT | DeepAccess | True | True |
| ACTCTGT-TGTAACTG | KMAC | False | N/A |
| TCTGTGTGTGT | KMAC | True | True |
| TGAGGACCGG | DeepAccess | True | False |
| ATGTTTAC-GTCACCTGAG | DeepAccess | True | True |
| CCTCCGCACC-GTAGGTGTTC | DeepAccess | True | True |
| TGATGTCACT | DeepAccess | True | True |
| ACGAATTATC-GTGACCACTG | DeepAccess | True | True |
| TATTGAC | KMAC | False | N/A |
| CCAGCACCCA-GTAGGTGTTC | DeepAccess | True | False |
| ATGTTTAC-GTGACCACTG | DeepAccess | True | True |
| TGCAGTGACA | DeepAccess | True | True |
| GAGTGACACT | DeepAccess | True | True |
| TGTAACTG | KMAC | False | N/A |
| TCCAGTTCCA | KMAC | False | N/A |
| GGTGTGGCC | KMAC | True | False |
| CCTCCGCACC | DeepAccess | True | True |
| GCCTTGTTGGAG | KMAC | False | N/A |
| ACTGGAACTC-TGCAAATTGT | KMAC | True | False |
| TATTGAC-TCCAGTTCCA | KMAC | False | N/A |
| GGAGTGACAC | DeepAccess | True | True |
| GCGCACACCT | DeepAccess | True | True |
| TATTGAC-GGCCATTGT | KMAC | False | N/A |
| CCTCCGCACC-GAACAGGTGC | DeepAccess | True | True |
| GTGACCACTG | DeepAccess | True | True |
| GTACCTGTTG | DeepAccess | True | False |
| ATACCGATAC | DeepAccess | False | N/A |
| GTCACCTGAG-GTGACCACTG | DeepAccess | True | True |
| ATGTAAACAT | KMAC | False | N/A |
| TCTGTGTGTGT-ACTGGAACTC | KMAC | False | N/A |
| ACGAATTATC-CGCGGCGGCC | DeepAccess | True | False |
| TGTTTTTGTT | KMAC | False | N/A |
| GTAGGTGTTC | DeepAccess | True | False |
| ACGAATTATC | DeepAccess | False | N/A |
| ATGTTTAC | DeepAccess | False | N/A |
| ATGTTTAC-ACGAATTATC | DeepAccess | False | N/A |
| CCAGCACCCA-CGCACACCTG | DeepAccess | False | N/A |
| GATTGGTTAA-ATACCGATAC | DeepAccess | False | N/A |
| GTCACCTGAG-CGCGGCGGCC | DeepAccess | False | N/A |
| CGCGGCGGCC-GTGACCACTG | DeepAccess | False | N/A |
| CAGCTGTG | KMAC | False | N/A |
| CTGCCTG | KMAC | False | N/A |
| TGTTTTTGTT-TGCAAATTGT | KMAC | True | False |
| GAACAGGTGC | DeepAccess | False | N/A |
| TGTTTTTGTT-CACCAGGGG | KMAC | False | N/A |
| CCTCCGCACC-GTACCTGTTG | DeepAccess | False | N/A |
| CCTCCGCACC-TGAGGACCGG | DeepAccess | True | True |
| TGTCACCTGA | DeepAccess | True | True |
| TGCAAATTGT | KMAC | False | N/A |
| CGCACACCTG | DeepAccess | True | True |
| TGCAGTGACA-TGATGTCACT | DeepAccess | True | True |
| ACTCTGT | KMAC | False | N/A |
| TGTCAACATT | DeepAccess | True | True |
| ACTGGAACTC | KMAC | False | N/A |
| ATACCGATAC-TGTTGTACCG | DeepAccess | True | True |
| TCTGTGTGTGT-CAGCTGTG | KMAC | True | True |
| GCGCACACCT-TGTTGTACCG | DeepAccess | True | True |
| TGATGTCACT-TGTCACCTGA | DeepAccess | True | True |
| CCAGCACCCA | DeepAccess | True | False |

Table S1. List of 76 motifs and motif pairs tested from DNase-seq motif discovery with KMAC and DeepAccess. Reported motifs that successfully drive differential accessibility between stem cell and definitive endoderm, and whether those that tested differential were also predicted to be differential from DeepAccess.

­­­

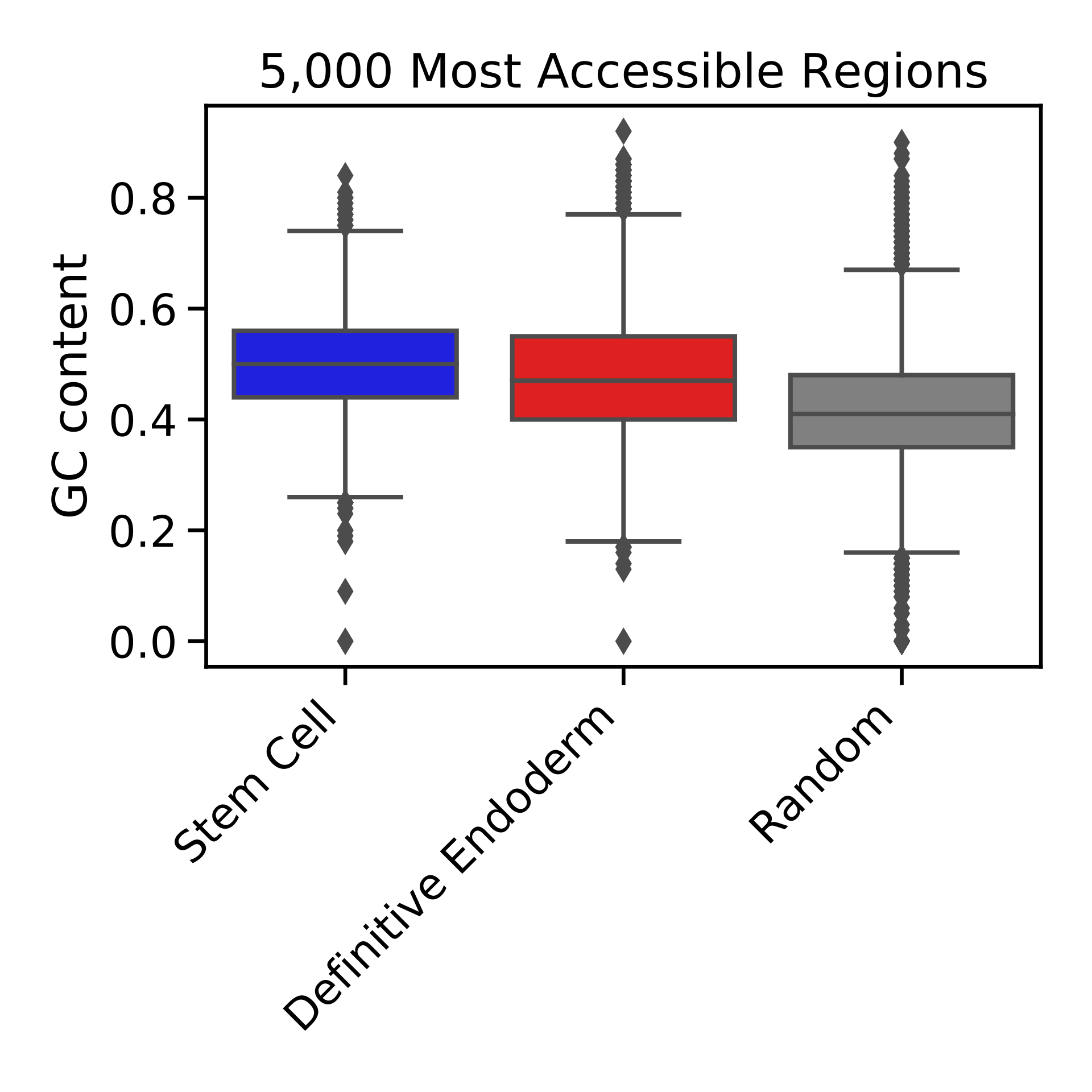

Figure S5. GC-content of top 5,000 most accessible genomic regions in stem cell and definitive endoderm is higher than in randomly selected closed regions (p < 0.001 by rank sum test for stem cell to random and p < 0.0001 by rank sum test definitive endoderm to random).

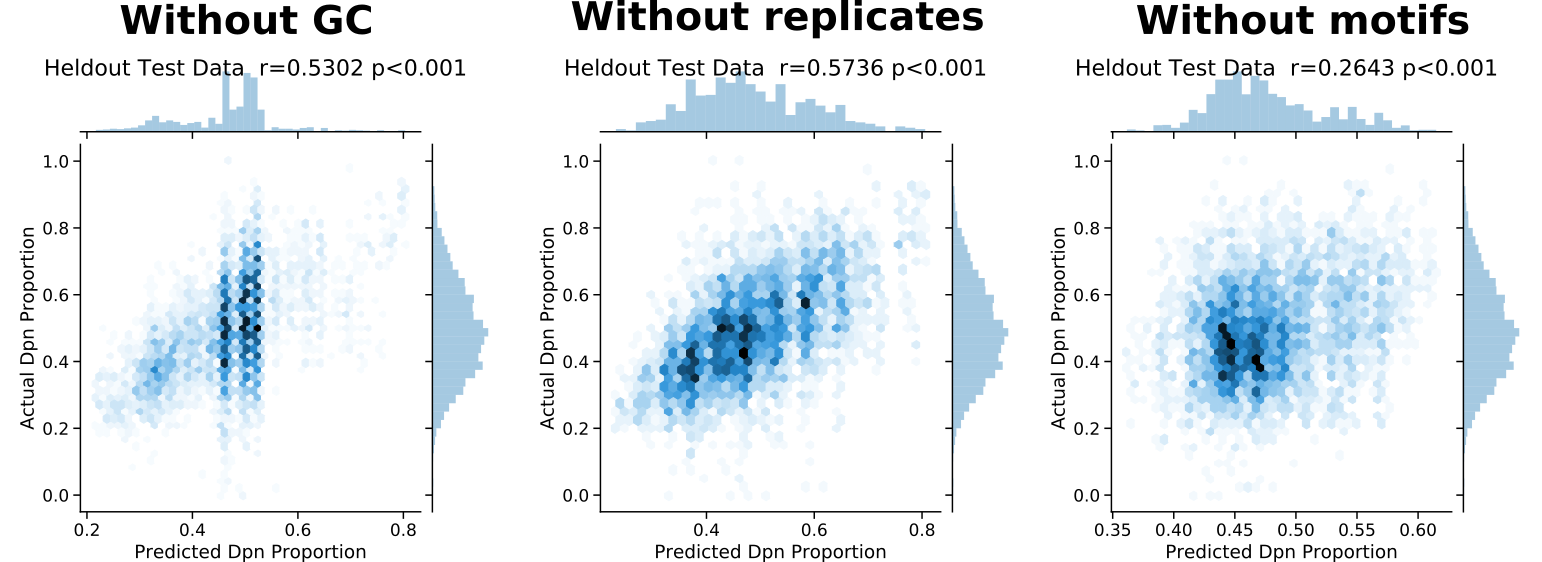

Figure S6. Contribution of modeling components of model trained on stem cell and definitive endoderm samples to Dpn proportion prediction shows motifs are a significant predictive component. Left panel, model performance on held out test data, trained without GC parameter. Center panel, model performance on held out test data, trained without replicate offset. Right panel, model performance on held out test data, trained without motifs. The correlation reported is the Pearson correlation coefficient (r).

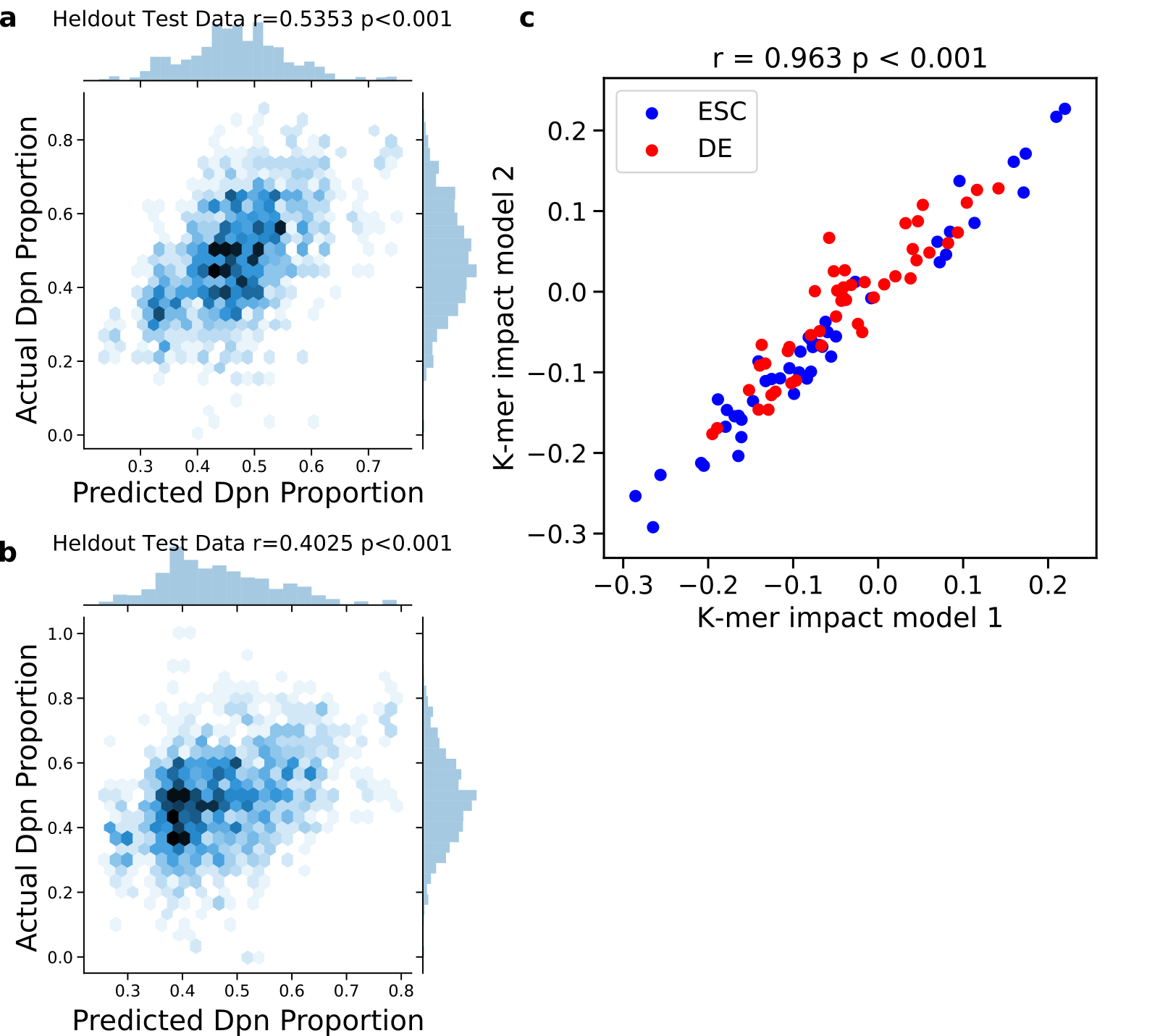

Figure S7. Regression model weights reproducible across biological replicates. We trained 2 regression models with features of GC-content, replicate, and cell type specific motifs. Model 1 is trained on experiments (es1.1, es1.2, ed1.1, ed1.2) and Model 2 is trained on experiments (es2.1, es2.2, ed2.1, ed2.2). A) Performance of Model 1 on held out test data. B) Performance of Model 2 on held out test data. C) Estimated cell type-specific motif weights of 2 regression models show high correlation. All reported R values are Pearson correlation coefficients.

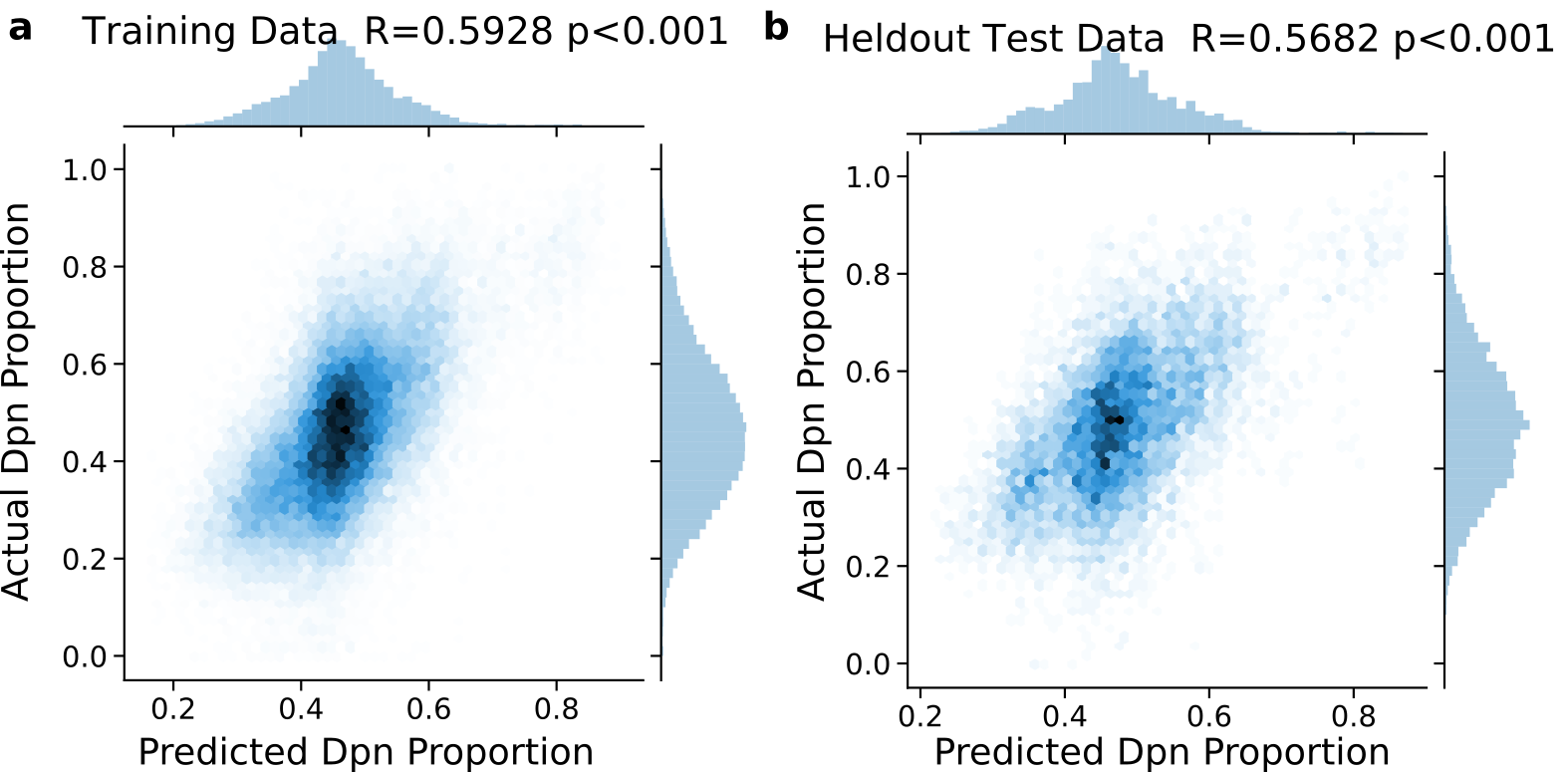

Figure S8. Training (n=34,528) and held-out (n=7,280) predictions of model trained on stem cell, definitive endoderm, Brachyury over-expression, and FoxA2 over-expression. The correlation reported is the Pearson correlation coefficient (r).

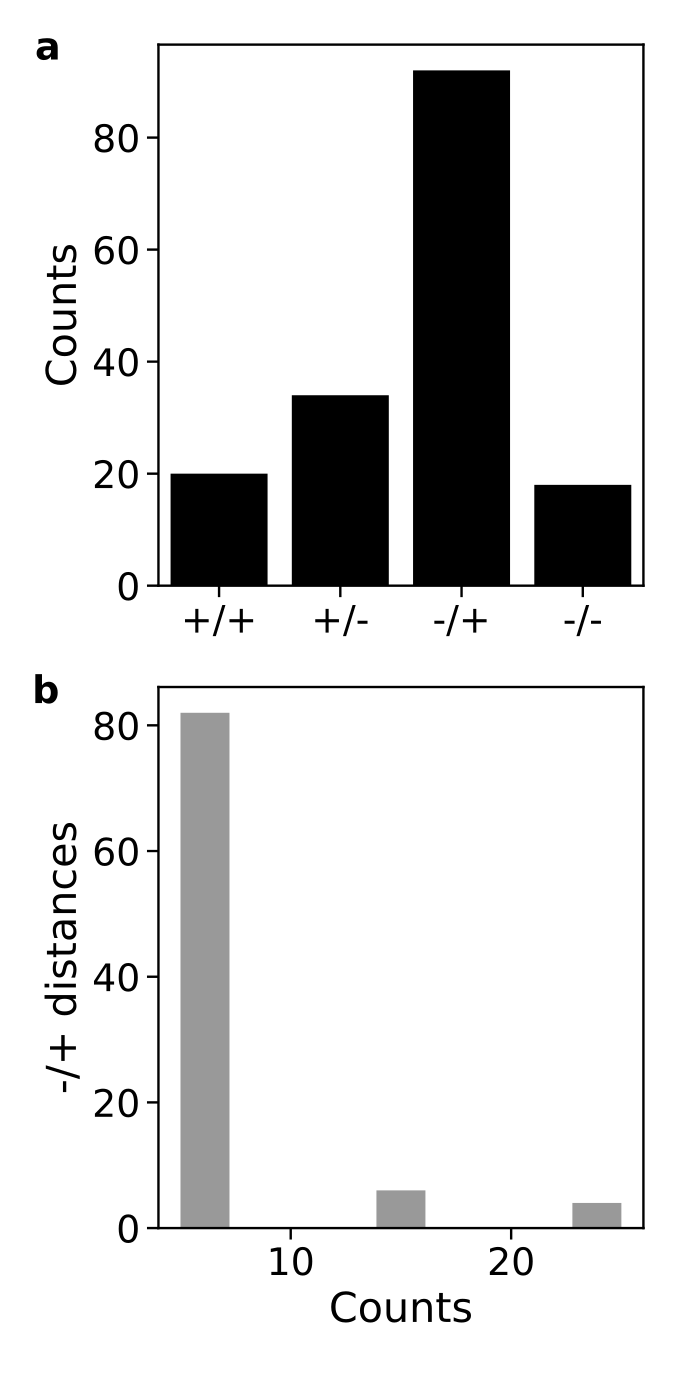

Figure S9. Brachyury dimers orientation and distance enrichment. A) Counts of Brachyury dimer motif orientations in Brachyury ChIP-seq peaks show statistical enrichment of motif pairs in ‘-/+’ orientation (p-value < 0.001 by Chi-squared test). B) Majority of ‘-/+’ motif instances (80/92) are 5 bp apart, indicating a spatial preference from dimerization.

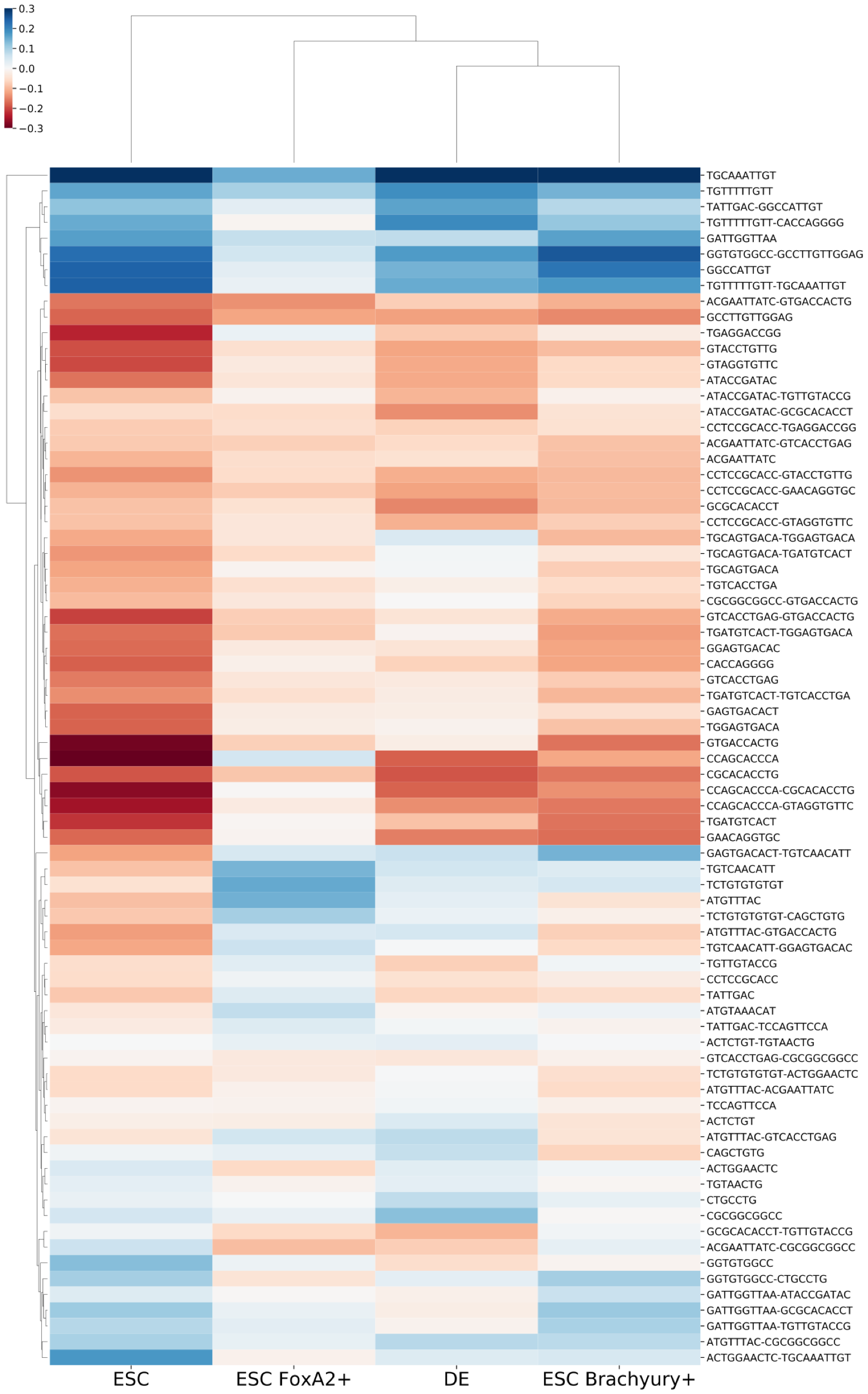
­­­

Figure S10. Full regression weights for all motifs for stem cells, definitive endoderm, FoxA2 over-expression, and Brachyury over-expression.

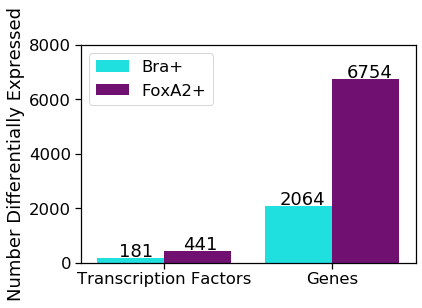

Figure S11. Differential expression analysis from over-expression of Brachyury and FoxA2 in Brachyury/Eomes knock-out definitive endoderm cells supports that more transcription changes occur due to FoxA2 expression.

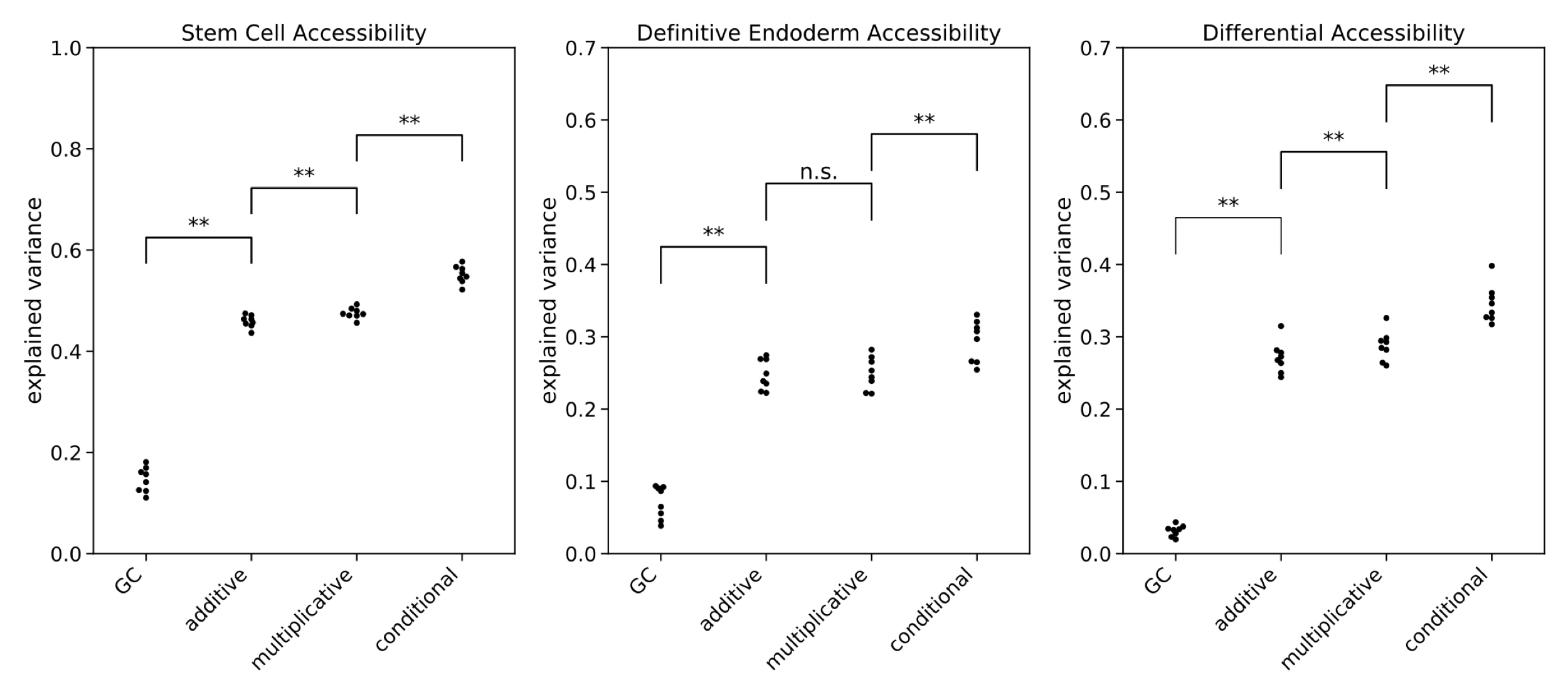

Figure S12. Comparing motif interaction models with conditional interactions between motif pairs compared to multiplicative (log additive), additive, and GC-content only regression model trained on average Dpn proportion to predict in stem cells and endoderm. Explained variance computed on held out test data (n = 1134-1209). Each dot is one of 8-fold trials computed on a sequence using a set of randomly selected 16/24 backgrounds as the training set (n = 2335-2413) and evaluated on test data from the other 8 backgrounds. A conditional interaction model has highest explained variance indicating presence of nonlinear interactions. P-values computed by one-tailed Wilcoxon rank sum test.

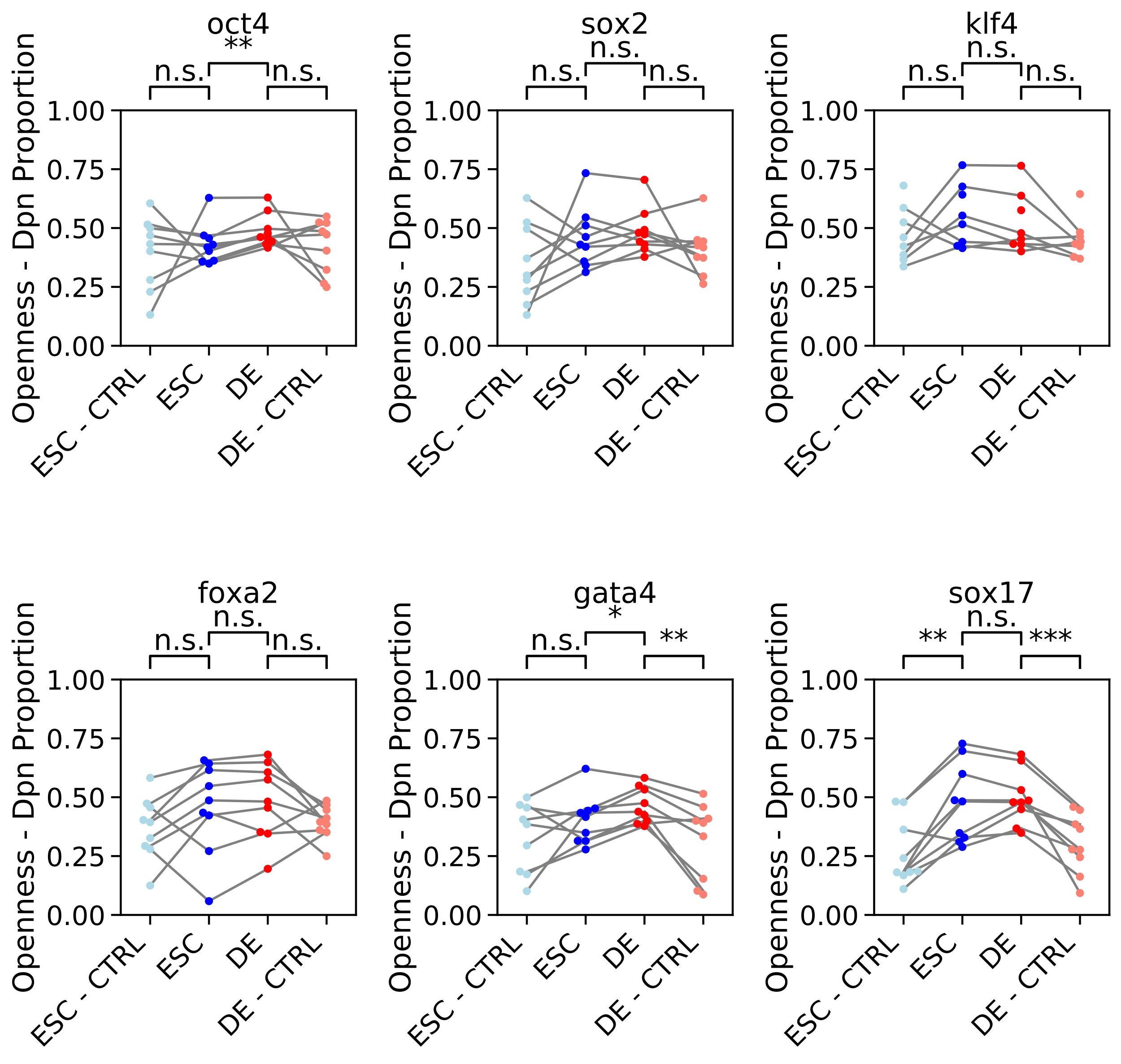

Figure S13. MIAA accessibility from single instances of pioneer transcription factor motifs. Single consensus motif instances of pioneer transcription factors inserted into neutral sequence backgrounds generally do not drive differential accessibility except Gata4, which is insignificant when adjusted for multiple hypothesis testing. Oct4 drives differential accessibility and surprisingly opens endoderm, but does not drive differential opening when compared to shuffled controls. All significance scores reported are unadjusted paired t-test.

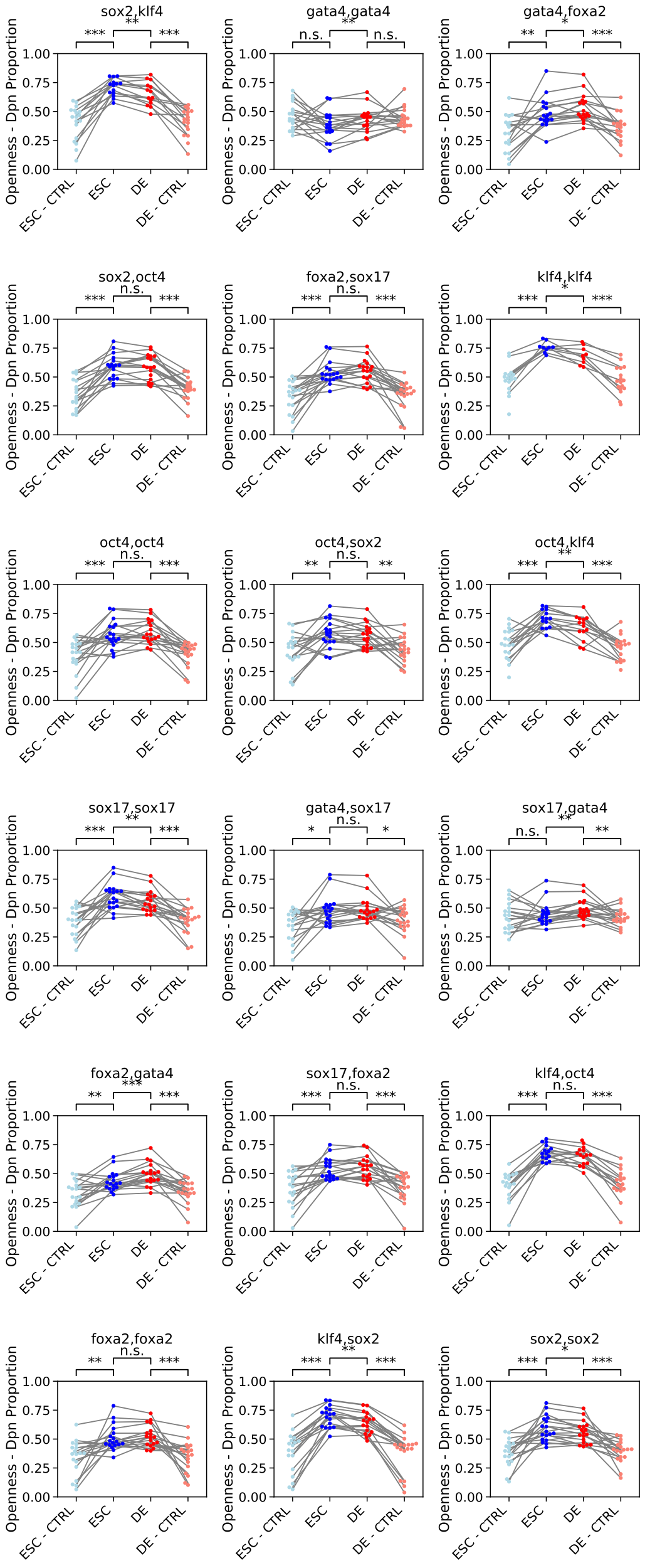

Figure S14. Phrases containing two motif instances are able to open chromatin in 17/18 cases (p < 0.05; Benjamini Hochberg multiple hypothesis correction) and differentially open chromatin between stem cell and endoderm in 9/18 cases (p < 0.05; Benjamini Hochberg multiple hypothesis correction). All significance values reported are unadjusted paired t-test.

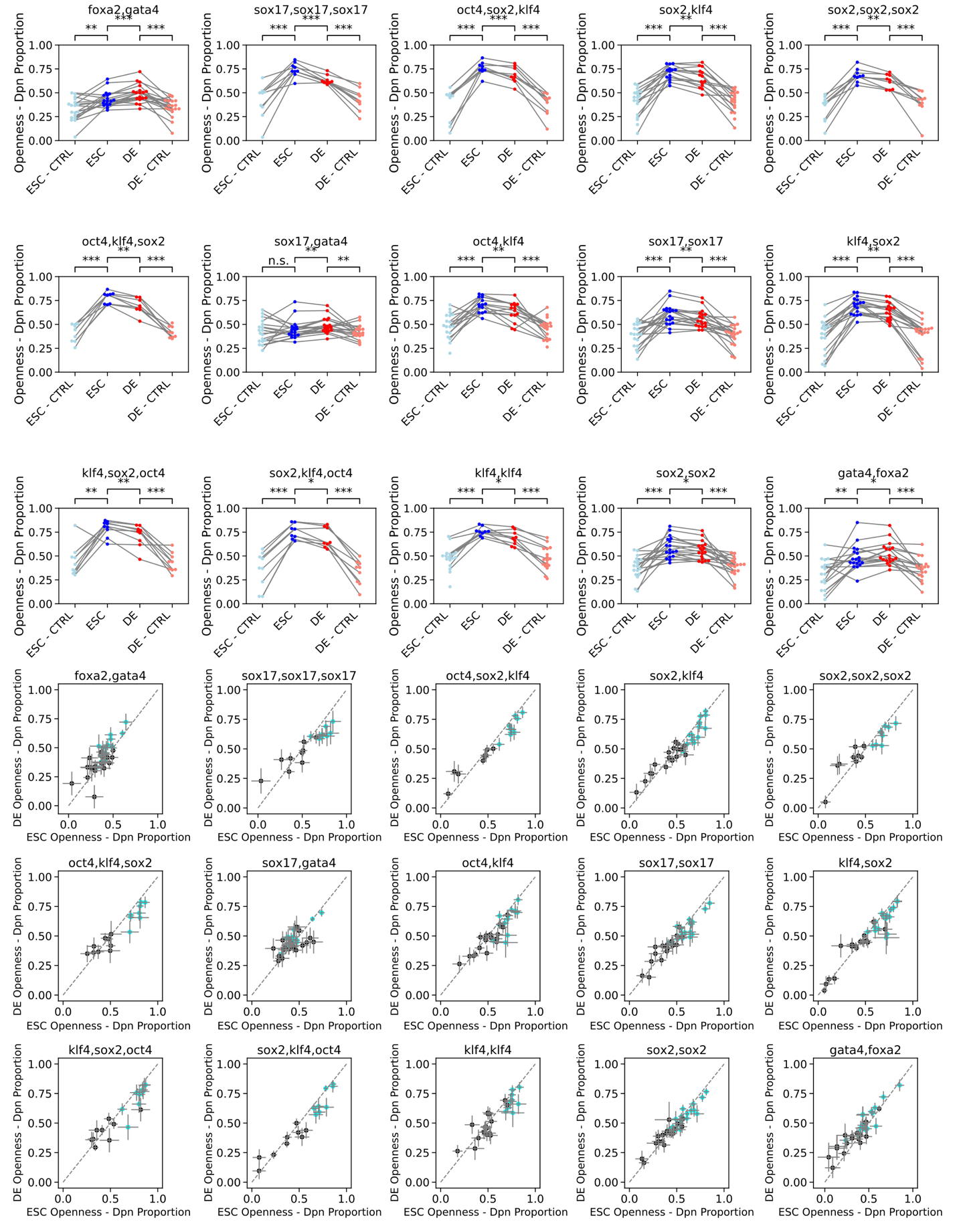

Figure S15. All differentially accessible pioneer motifs or combinations of pioneer motifs. 15 out of 42 possible orderings and combinations of 1, 2, or 3 motif sites are differentially accessible under multiple hypothesis testing. Top shows dot plot of MIAA measured accessibility of each phrase over 9 (1 or 3 motif instances) or 18 (2 motif instances, measuring 2 distances) different sequence backgrounds, and compared to randomly perturbed control sequences. Bottom shows each phrase’s MIAA measured accessibility in stem cell (ESC) and definitive endoderm (DE) along with 95% confidence intervals from experimental replicates of MIAA assay.

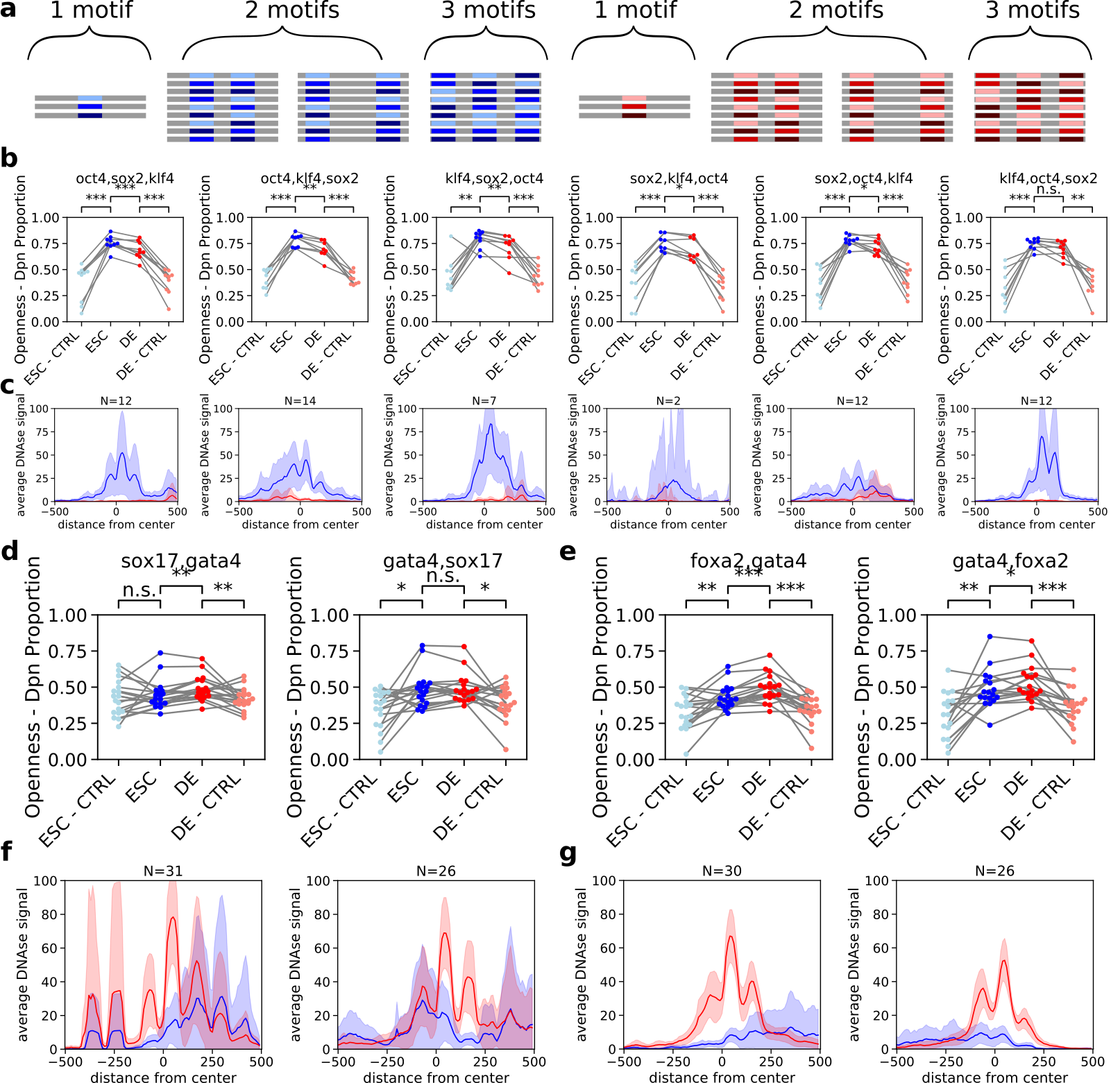

Figure S16. MIAA and DNase-seq reflect ordering-based logic of pioneer transcription factor action. A) Single and combinations of Oct4, Sox2, and Klf4 (left, ESC pioneers) and FoxA2, Gata4, and Sox17 (right, DE pioneers) generated with all possible orders of motifs with 1, 2 (distance of 6 and 20 apart), or 3 (distance of 6 apart) instances of motifs. B) Ordering of 3 stem cell pioneers Oct4, Sox2, and Klf4 results in different levels of differential accessibility, which is also reflected by average DNase-seq profiles and 95% confidence intervals of genomic sites containing ordered motifs. C) Sox17 and Gata4 differentially accessible when Sox17 is first in pair and not when Sox17 second in pair. D) Foxa2 and Gata4 more differentially accessible when Foxa2 is first in pair and less when Foxa2 is second in pair. All p-values reported are uncorrected paired t-test.

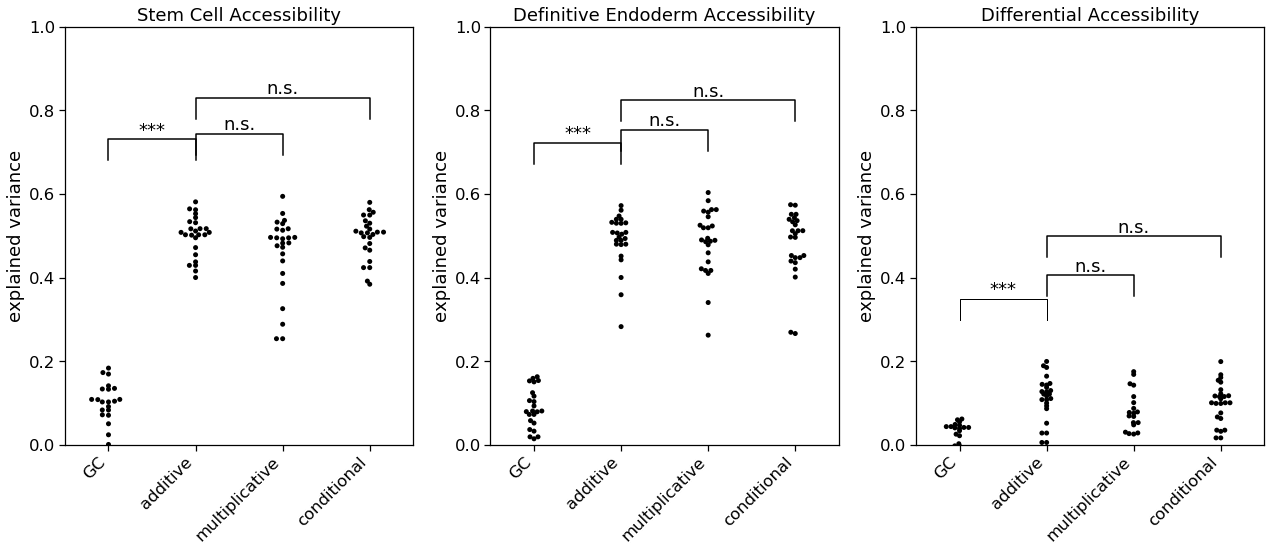

Figure S17. Comparison of regression models on phrase library of combinations of transcription factors. No significant difference between additive, multiplicative, or conditional models for predicting accessibility for stem cell, definitive endoderm, or differential accessibility. P-values computed by one-tailed Wilcoxon Signed-Rank test.

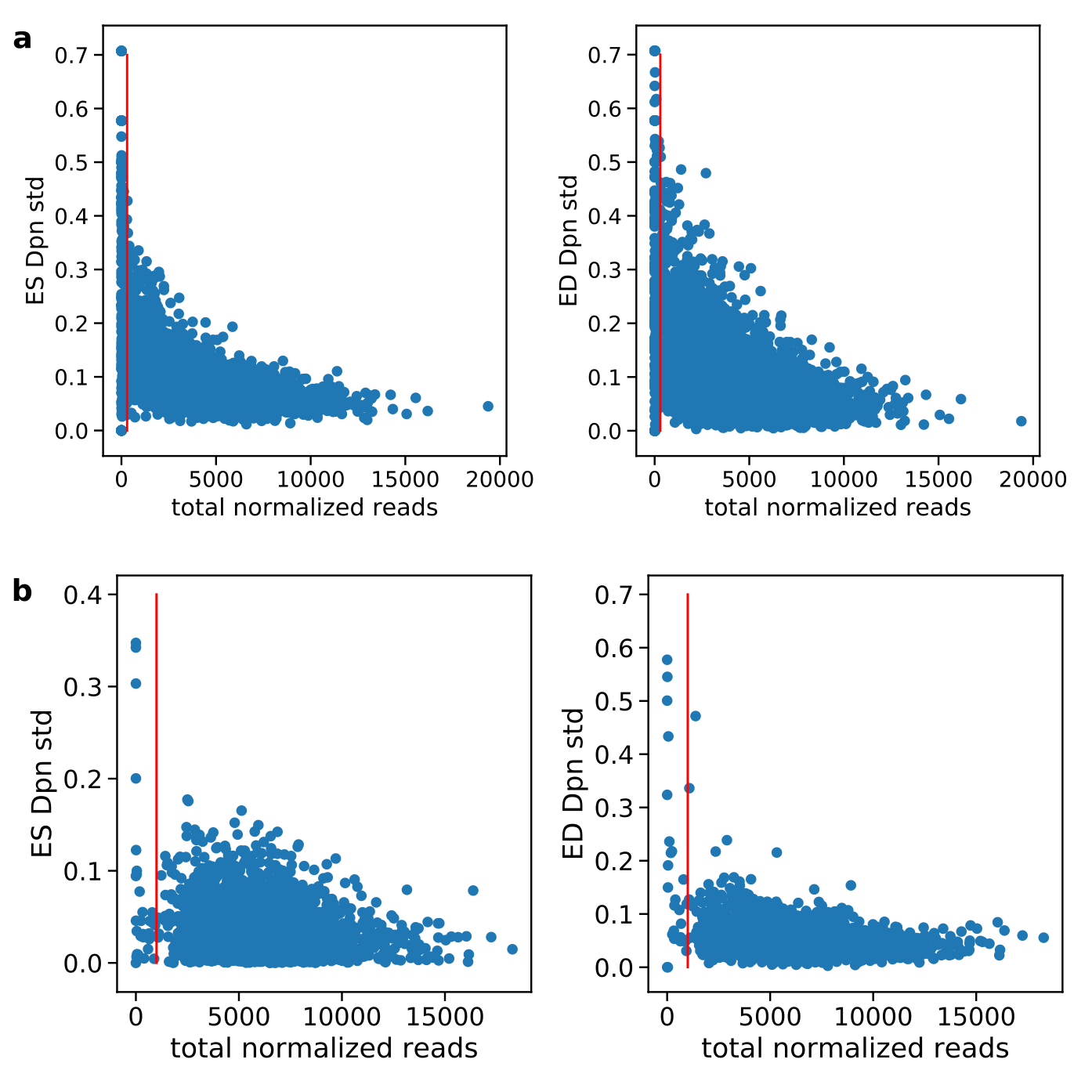

Figure S18. A) 6,000 phrase library designed using motifs from DNase-seq enrichment. Each dot represents a phrase measured across multiple replicates. Y-axis represents Dpn proportion (MIAA-measured openness) over replicates in stem cells (ES) and definitive endoderm (ED). Red line represents threshold for selecting phrases with sufficient read coverage and lower standard deviations. B) 2,000 phrase library designed using pioneer transcription factor consensus motifs. Red line represents phrase threshold for selecting phrases with sufficient read coverage to have low standard deviation over experimental replicates.

|  | TF-combo | Best Model ESC | Best Model DE | Best Model Differential | Is Differential |
| --- | --- | --- | --- | --- | --- |
| 0 | sox2,oct4 | additive | additive | additive | False |
| 1 | oct4,sox2 | additive | multiplicative | additive | False |
| 2 | sox2,sox2 | additive | additive | additive | True |
| 3 | klf4,sox2 | additive | multiplicative | additive | True |
| 4 | sox2,klf4 | additive | additive | additive | True |
| 5 | oct4,klf4 | additive | additive | additive | True |
| 6 | klf4,klf4 | additive | additive | additive | True |
| 7 | oct4,oct4 | additive | additive | additive | False |
| 8 | klf4,oct4 | additive | additive | additive | False |
| 9 | klf4,oct4,sox2 | additive | additive | additive | False |
| 10 | klf4,sox2,oct4 | additive | additive | additive | True |
| 11 | sox2,klf4,oct4 | additive | additive | additive | True |
| 12 | oct4,oct4,oct4 | additive | additive | additive | False |
| 13 | sox2,oct4,klf4 | additive | additive | additive | False |
| 14 | sox2,sox2,sox2 | additive | additive | additive | True |
| 15 | oct4,sox2,klf4 | additive | additive | additive | True |
| 16 | oct4,klf4,sox2 | additive | additive | additive | True |
| 17 | klf4,klf4,klf4 | additive | additive | conditional | False |
| 18 | sox17,sox17 | additive | additive | additive | True |
| 19 | foxa2,sox17 | additive | additive | additive | False |
| 20 | gata4,gata4 | multiplicative | additive | additive | False |
| 21 | gata4,sox17 | additive | additive | additive | False |
| 22 | sox17,foxa2 | additive | additive | additive | False |
| 23 | sox17,gata4 | multiplicative | additive | multiplicative | True |
| 24 | foxa2,gata4 | additive | additive | additive | True |
| 25 | gata4,foxa2 | additive | additive | additive | True |
| 26 | foxa2,foxa2 | additive | additive | additive | False |
| 27 | foxa2,sox17,gata4 | additive | multiplicative | additive | False |
| 28 | gata4,gata4,gata4 | multiplicative | additive | multiplicative | False |
| 29 | foxa2,foxa2,foxa2 | additive | multiplicative | additive | False |
| 30 | foxa2,gata4,sox17 | additive | additive | additive | False |
| 31 | sox17,foxa2,gata4 | additive | additive | additive | False |
| 32 | gata4,sox17,foxa2 | multiplicative | additive | multiplicative | False |
| 33 | sox17,sox17,sox17 | additive | additive | additive | True |
| 34 | sox17,gata4,foxa2 | additive | additive | additive | False |
| 35 | gata4,foxa2,sox17 | additive | additive | additive | False |

Table S2. Best models for each transcription factor combination, selected by significant decrease (p<0.05 by Wilcoxon Signed-Rank Test) in mean-squared error over 25-fold cross validation. Last column indicates transcription factor combinations that were called differentially accessible between cell types and compared to shuffled control phrases. ­­­

| Description | Data Type | Accession |
| --- | --- | --- |
| Brachyury/Eomes Knock-out mouse ES cells w/ overexpression FoxA2 or Brachyury | RNA-seq | GSE128466 |
| Brachyury ChIP-seq mesendoderm | ChIP-seq | GSE54978 |
| FoxA2 ChIP-seq definitive endoderm | ChIP-seq | GSE116258 |
| DNase-seq stem cell and definitive endoderm | DNase-seq | GSE53776 |

Table S3. Accession of public data used in analysis.
